## Supplemental Figure S1 for "*Ciona Brachyury* proximal and distal enhancers have different FGF dose-response relationships"

**Supplemental Figure 1. Parameter distributions from bootstrapped residuals.**

**(A-C)** Bootstrap parameter distributions for curves fit to the **(A)** Proximal, **(B)** Distal, and **(C)** Full Length whole-embryo reporter data. Monophasic Hill functions are fit for the Proximal and Distal constructs. A biphasic double-sigmoid function is fit for the Full Length construct.

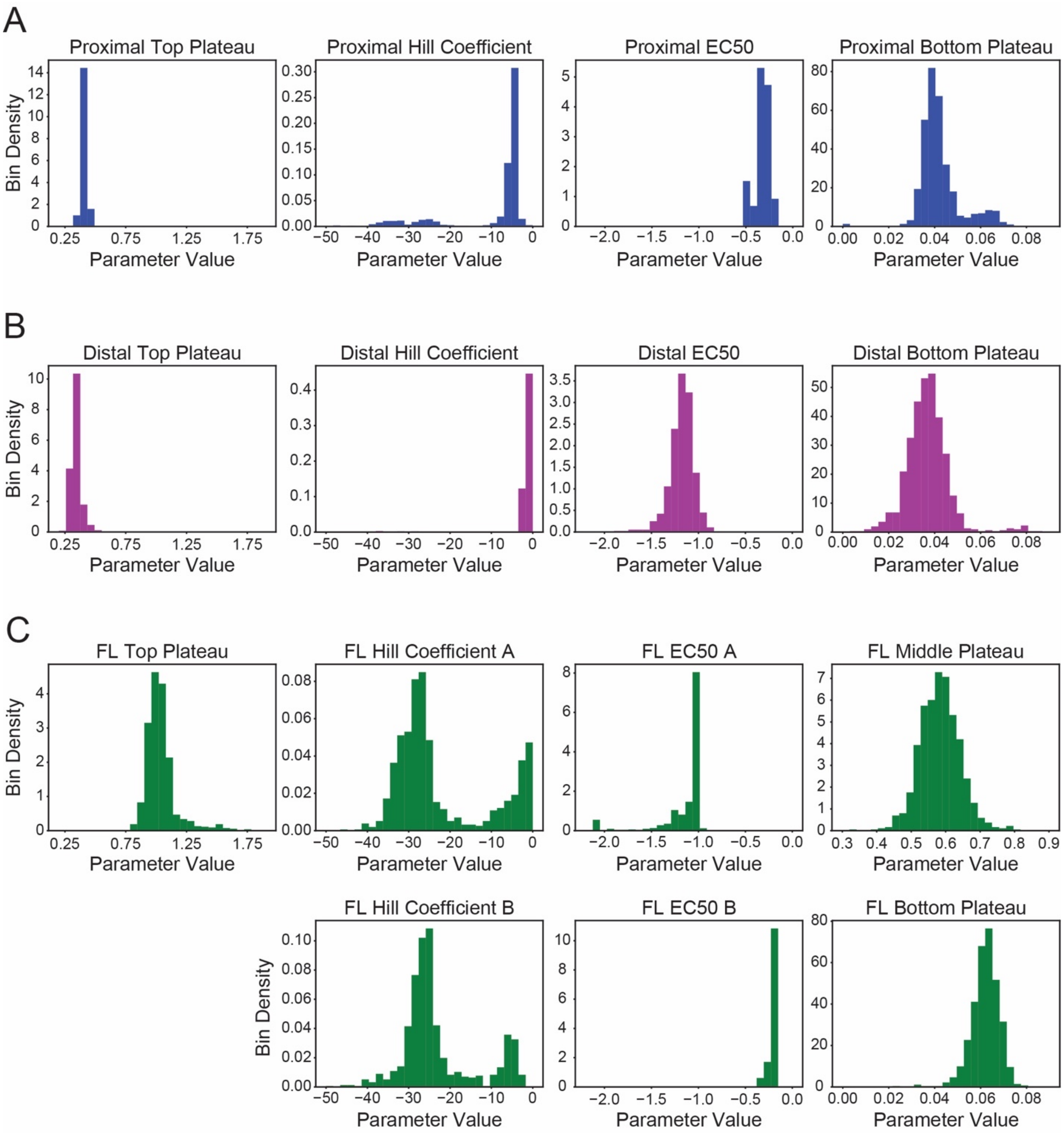
