## Supplemental Appendix S3 for "*Ciona Brachyury* proximal and distal enhancers have different FGF dose-response relationships": Fig 2 and Associated Supplement.py

```
#!/usr/bin/env python3
# -*- coding: utf-8 -*-
"""
Created on Wed May 20 16:06:43 2020

@author: Matt
"""
#Built to be the final repository of code for all things to do with the SEV plotting
#This is NOT a place to tinker with ideas or generate completely new plots
#ONLY use this to make adjustments for final versions of figure panels, and as a script to produce them

#Import necessary libraries
import pandas as pd
import os
import matplotlib.pyplot as plt
import seaborn as sns
plt.rcParams['pdf.fonttype']=42
import matplotlib.lines as mlines
import numpy as np
from scipy.optimize import curve_fit
import math
from scipy.stats import ttest_ind
from scipy.stats import ttest_ind_from_stats
from scipy.stats import ranksums

#Change directory for input/output of data, read in and split data into useful subdivisions
os.chdir("/Volumes/MH Veeman Lab Home Backup/SEV Plots and Data")
SEVScores=pd.read_csv('SEVData.csv')
Prox=SEVScores[SEVScores['enhancer_construct']=="Proximal"]
ProxDoses=Prox['ModLogDose']
ProxScores=Prox['NSSEVNormedToFL0.01581']
PStoAdd=ProxScores.tolist()
Shadow=SEVScores[SEVScores['enhancer_construct']=="Shadow"]
ShadowDoses=Shadow['ModLogDose']
ShadowScores=Shadow['NSSEVNormedToFL0.01581']
SStoAdd=ShadowScores.tolist()
Full=SEVScores[SEVScores['enhancer_construct']=="Shadow+Spacer+Proximal"]
FullDoses=Full['ModLogDose']
FullScores=Full['NSSEVNormedToFL0.01581']
FStoAdd=FullScores.tolist()
detailedx=np.linspace(-2.1,0.61,100)
ProxN=len(Prox)
ShadowN=len(Shadow)
FullN=len(Full)

#Enter plotting settings that will be referenced repeatedly
doseticks=[np.log10(.00855),np.log10(.02924),np.log10(0.05408),np.log10(.1),np.log10(.1849),np.log10(.342),np.log10(.6325),np.log10(1.17),np.log10(2.163),np.log10(4)]
doselabels=['DMSO','0.029','0.054','0.10','0.18','0.34','0.63','1.2','2.2','4.0']

#Make line plot to show data from all reporters/doses on the same axis
pline=mlines.Line2D([],[],color='b',label='Proximal')
dline=mlines.Line2D([],[],color='m',label='Distal')
fline=mlines.Line2D([],[],color='g',label='Full Length')
fig,ax=plt.subplots(figsize=(8,6))
fig=sns.lineplot(x="ModLogDose",y="NSSEVNormedToFL0.01581",hue="enhancer_construct",palette=["blue","green","magenta"],markers=['o','o','o'],dashes=False,style='enhancer_construct',ax=ax,ci=95,data=SEVScores)
fig.set(xlabel="[U0126 (uM)]",xticks=doseticks,xticklabels=doselabels,ylabel="Normalized Expression",title='SEV by Reporter')
plt.setp(ax,ylim=(-0.02,1.5))
fig=plt.legend(handles=[pline,dline,fline])
fig.figure.savefig('FINAL VERSION SEV Plot.pdf',transparent=True)

#Address the question of additivity at the DMSO dose by bootstrapping
Additive=SEVScores[SEVScores['perturbation_name']==0.01581]
ProxC=Additive[Additive['enhancer_construct']=='Proximal']
ProxC=ProxC['NSSEVNormedToFL0.01581'].tolist()
DistalC=Additive[Additive['enhancer_construct']=='Shadow']
DistalC=DistalC['NSSEVNormedToFL0.01581'].tolist()
FLC=Additive[Additive['enhancer_construct']=='Shadow+Spacer+Proximal']
FLC=FLC['NSSEVNormedToFL0.01581'].tolist()

ColumnNames=[]
for x in range(1000):
    iteration=x+1
    name='iter'+str(iteration)
    ColumnNames.append(name)

ProxBSC=np.random.choice(ProxC,(len(ProxC),1000))
ProxBSC=pd.DataFrame(ProxBSC,columns=ColumnNames)
DistalBSC=np.random.choice(DistalC,(len(DistalC),1000))
DistalBSC=pd.DataFrame(DistalBSC,columns=ColumnNames)
FLBSC=np.random.choice(FLC,(len(FLC),1000))
FLBSC=pd.DataFrame(FLBSC,columns=ColumnNames)
ProxBSC.to_csv('FINAL VERSION Proximal Control Embryo SEVs Bootstrapped.csv')
DistalBSC.to_csv('FINAL VERSION Distal Control Embryo SEVs Bootstrapped.csv')
FLBSC.to_csv('FINAL VERSION FL Control Embryo SEVs Bootstrapped.csv')

#Obtain means of each bootstrap, store in a dataframe for use in stats and plotting
PMeans=[]
DMeans=[]
FMeans=[]
SumMeans=[]
for column in ColumnNames:
    pmean=np.average(ProxBSC[column])
    dmean=np.average(DistalBSC[column])
    fmean=np.average(FLBSC[column])
    summean=pmean+dmean
    PMeans.append(pmean)
    DMeans.append(dmean)
    FMeans.append(fmean)
    SumMeans.append(summean)
BSAdd=pd.DataFrame()
BSAdd['BSRep']=ColumnNames
BSAdd['Proximal Means']=PMeans
BSAdd['Distal Means']=DMeans
BSAdd['FL Means']=FMeans
BSAdd['Sum of P and D Means']=SumMeans

#Make a barplot of the bootstrapped reporter at DMSO
bstoplot=[np.average(BSAdd['Proximal Means']),np.average(BSAdd['Distal Means']),np.average(BSAdd['Sum of P and D Means']),np.average(BSAdd['FL Means'])]
bspl,bspu=np.percentile(BSAdd['Proximal Means'],[2.5,97.5])
bsdl,bsdu=np.percentile(BSAdd['Distal Means'],[2.5,97.5])
bssl,bssu=np.percentile(BSAdd['Sum of P and D Means'],[2.5,97.5])
bsfl,bsfu=np.percentile(BSAdd['FL Means'],[2.5,97.5])
bsplotcolors=['b','m','c','g']
fig,ax=plt.subplots(figsize=(8,6))
fig=plt.bar([1,2,3,4],bstoplot,color=bsplotcolors,yerr=[[bstoplot[0]-bspl,bstoplot[1]-bsdl,bstoplot[2]-bssl,bstoplot[3]-bsfl],[bspu-bstoplot[0],bsdu-bstoplot[1],bssu-bstoplot[2],bsfu-bstoplot[3]]])
plt.setp(ax,xlabel='Reporter',xticks=[1,2,3,4],xticklabels=['Proximal','Distal','Sum','Full Length'],ylabel='Normalized Expression')
plt.savefig('FINAL VERSION Bootstrapped Additivity Question.pdf',transparent=True)

#Run t-tests to compare Proximal and Distal, as well as Sum v FL
stat,p=ttest_ind(BSAdd['Sum of P and D Means'],BSAdd['FL Means'])
pdstat,pdp=ttest_ind(BSAdd['Proximal Means'],BSAdd['Distal Means'])
AdditivityResults=pd.DataFrame()
AdditivityResults['Comparison']=['SumVFL','PvD']
AdditivityResults['Statistic']=[stat,pdstat]
AdditivityResults['P-value']=[p,pdp]
AdditivityResults.to_csv('FINAL VERSION Additivity at DMSO.csv')

#Add curve fitting programs, with bounds defined as the least restrictive bounds possible
#For linear curves
def linefit(x,M,B):
    return M*x+B
def residualsline(p,y,x):
    M,B=p
    err=y-linefit(x,M,B)
    return err
def pevalline(x,p):
    M,B=p
    return linefit(x,M,B)
BoundsLine=([-np.inf,0],[0,np.inf])

#For 4-parameter monophasic curves
def logistic4(x,A,B,C,D):
    return ((A-D)/(1+(10**((C-x)*B))))+D
def residuals4(p,y,x):
    A,B,C,D=p
    err=y-logistic4(x,A,B,C,D)
    return err
def peval4(x,p):
    A,B,C,D=p
    return logistic4(x,A,B,C,D)
Bounds4=([0.2,-np.inf,-np.inf,0],[np.inf,0,np.inf,0.2])

#For 7-parameter biphasic curves
def logistic7(x,A,B,C,D,E,F,G):
    return ((A-D)/(1.0+(10**((C-x)*B))))+((D-G)/(1.0+(10**((F-x)*E))))+G
def residuals7(p,y,x):
    A,B,C,D,E,F,G=p
    err=y-logistic7(x,A,B,C,D,E,F,G)
    return err
def peval7(x,p):
    A,B,C,D,E,F,G=p
    return logistic7(x,A,B,C,D,E,F,G)
Bounds7=([0.2,-np.inf,np.log10(0.00855),-np.inf,-np.inf,np.log10(0.3420),0],[np.inf,0,np.log10(0.1849),np.inf,0,np.log10(4),0.2])

#Fit curves of each type for each reporter, using unbiased guesses, to determine which type is best for the particular reporter
ProxLinOpt,ProxLinCov=curve_fit(linefit,ProxDoses,ProxScores,bounds=BoundsLine)
Prox4Opt,Prox4Cov=curve_fit(logistic4,ProxDoses,ProxScores,bounds=Bounds4)
Prox7Opt,Prox7Cov=curve_fit(logistic7,ProxDoses,ProxScores,bounds=([0.2,-np.inf,-np.inf,-np.inf,-np.inf,-np.inf,0],[np.inf,0,np.inf,np.inf,0,np.inf,0.2]))
ProxLinErr=np.diagonal(ProxLinCov)
Prox4Err=np.diagonal(Prox4Cov)
Prox7Err=np.diagonal(Prox7Cov)
ProxLinRes=residualsline(ProxLinOpt,ProxScores,ProxDoses)
Prox4Res=residuals4(Prox4Opt,ProxScores,ProxDoses)
Prox7Res=residuals7(Prox7Opt,ProxScores,ProxDoses)

ShadowLinOpt,ShadowLinCov=curve_fit(linefit,ShadowDoses,ShadowScores,bounds=BoundsLine)
Shadow4Opt,Shadow4Cov=curve_fit(logistic4,ShadowDoses,ShadowScores,bounds=Bounds4)
Shadow7Opt,Shadow7Cov=curve_fit(logistic7,ShadowDoses,ShadowScores,bounds=Bounds7)
ShadowLinErr=np.diagonal(ShadowLinCov)
Shadow4Err=np.diagonal(Shadow4Cov)
Shadow7Err=np.diagonal(Shadow7Cov)
ShadowLinRes=residualsline(ShadowLinOpt,ShadowScores,ShadowDoses)
Shadow4Res=residuals4(Shadow4Opt,ShadowScores,ShadowDoses)
Shadow7Res=residuals7(Shadow7Opt,ShadowScores,ShadowDoses)

FullLinOpt,FullLinCov=curve_fit(linefit,FullDoses,FullScores,bounds=BoundsLine)
Full4Opt,Full4Cov=curve_fit(logistic4,FullDoses,FullScores,bounds=Bounds4)
Full7Opt,Full7Cov=curve_fit(logistic7,FullDoses,FullScores,bounds=Bounds7)
FullLinErr=np.diagonal(FullLinCov)
Full4Err=np.diagonal(Full4Cov)
Full7Err=np.diagonal(Full7Cov)
FullLinRes=residualsline(FullLinOpt,FullScores,FullDoses)
Full4Res=residuals4(Full4Opt,FullScores,FullDoses)
Full7Res=residuals7(Full7Opt,FullScores,FullDoses)

#Plot out curve options for FL to show the different fit styles
linear=mlines.Line2D([],[],color='g',ls=':',label='Linear')
mono=mlines.Line2D([],[],color='g',ls='--',label='Monophasic')
bi=mlines.Line2D([],[],color='g',ls='-.',label='Biphasic')
fig,ax=plt.subplots(figsize=(8,6))
fig=sns.lineplot(x="ModLogDose",y="NSSEVNormedToFL0.01581",color='g',markers=['o'],dashes=False,style='enhancer_construct',ax=ax,ci=95,data=Full)
fig=plt.plot(detailedx,pevalline(detailedx,FullLinOpt),':g')
fig=plt.plot(detailedx,peval4(detailedx,Full4Opt),'--g')
fig=plt.plot(detailedx,peval7(detailedx,Full7Opt),'-.g')
fig=plt.xticks(ticks=doseticks,labels=doselabels)
fig=plt.xlabel("[U0126 (uM)]")
fig=plt.ylabel("Normalized Expression")
fig=plt.title('FL Curve Selection')
plt.setp(ax,ylim=(-0.02,1.5))
fig=plt.legend(handles=[fline,linear,mono,bi])
fig=plt.savefig('FINAL VERSION FL Curve Selection.pdf',transparent=True)

#Calculate AIC and relative likelihood values for each fit in order to highlight the best curve type for each reporter
ProxLinRSS=np.sum(ProxLinRes*ProxLinRes)
Prox4RSS=np.sum(Prox4Res*Prox4Res)
Prox7RSS=np.sum(Prox7Res*Prox7Res)
ShadowLinRSS=np.sum(ShadowLinRes*ShadowLinRes)
Shadow4RSS=np.sum(Shadow4Res*Shadow4Res)
Shadow7RSS=np.sum(Shadow7Res*Shadow7Res)
FullLinRSS=np.sum(FullLinRes*FullLinRes)
Full4RSS=np.sum(Full4Res*Full4Res)
Full7RSS=np.sum(Full7Res*Full7Res)

ProxLinAIC=ProxN*np.log(ProxLinRSS/ProxN)+(2*3)
Prox4AIC=ProxN*np.log(Prox4RSS/ProxN)+(2*5)
Prox7AIC=ProxN*np.log(Prox7RSS/ProxN)+(2*8)
ShadowLinAIC=ShadowN*np.log(ShadowLinRSS/ShadowN)+(2*3)
Shadow4AIC=ShadowN*np.log(Shadow4RSS/ShadowN)+(2*5)
Shadow7AIC=ShadowN*np.log(Shadow7RSS/ShadowN)+(2*8)
FullLinAIC=FullN*np.log(FullLinRSS/FullN)+(2*3)
Full4AIC=FullN*np.log(Full4RSS/FullN)+(2*5)
Full7AIC=FullN*np.log(Full7RSS/FullN)+(2*8)

MinProxAIC=min(ProxLinAIC,Prox4AIC,Prox7AIC)
MinShadowAIC=min(ShadowLinAIC,Shadow4AIC,Shadow7AIC)
MinFullAIC=min(FullLinAIC,Full4AIC,Full7AIC)

ProxLinRL=math.exp((MinProxAIC-ProxLinAIC)/2)
Prox4RL=math.exp((MinProxAIC-Prox4AIC)/2)
Prox7RL=math.exp((MinProxAIC-Prox7AIC)/2)
ShadowLinRL=math.exp((MinShadowAIC-ShadowLinAIC)/2)
Shadow4RL=math.exp((MinShadowAIC-Shadow4AIC)/2)
Shadow7RL=math.exp((MinShadowAIC-Shadow7AIC)/2)
FullLinRL=math.exp((MinFullAIC-FullLinAIC)/2)
Full4RL=math.exp((MinFullAIC-Full4AIC)/2)
Full7RL=math.exp((MinFullAIC-Full7AIC)/2)

ReporterList=['Proximal','Proximal','Proximal','Distal','Distal','Distal','Full Length','Full Length','Full Length']
CurveTypeList=['Linear','Monophasic','Biphasic','Linear','Monophasic','Biphasic','Linear','Monophasic','Biphasic']
AICList=[ProxLinAIC,Prox4AIC,Prox7AIC,ShadowLinAIC,Shadow4AIC,Shadow7AIC,FullLinAIC,Full4AIC,Full7AIC]
RLList=[ProxLinRL,Prox4RL,Prox7RL,ShadowLinRL,Shadow4RL,Shadow7RL,FullLinRL,Full4RL,Full7RL]
CurveSelection=pd.DataFrame(columns=['Reporter','CurveType','AIC','RelativeLikelihood'])
CurveSelection.iloc[:,0]=ReporterList
CurveSelection.iloc[:,1]=CurveTypeList
CurveSelection.iloc[:,2]=AICList
CurveSelection.iloc[:,3]=RLList
CurveSelection.to_csv("FINAL VERSION Curve Selection Table.csv")

#Bootstrap the residuals of the best curve type for each reporter, limiting bootstrapping to within doses
BigDoseList=ProxDoses
DoseList=[]
for dose in BigDoseList:
    if dose not in DoseList:
        DoseList.append(dose)

PSortedDoses=[]
PSortedScored=[]
PSortedRes=pd.DataFrame(columns=ColumnNames)
SSortedDoses=[]
SSortedScored=[]
SSortedRes=pd.DataFrame(columns=ColumnNames)
FSortedDoses=[]
FSortedScored=[]
FSortedRes=pd.DataFrame(columns=ColumnNames)
for dose in DoseList:
    pdosematch=ProxDoses[ProxDoses==dose]
    presmatch=Prox4Res[ProxDoses==dose]
    pscorematch=ProxScores[ProxDoses==dose].tolist()
    PSortedScored=PSortedScored+pscorematch
    pbsres=np.random.choice(presmatch,(len(presmatch),1000))
    pbsres=pd.DataFrame(data=pbsres,columns=ColumnNames)
    PSortedRes=pd.concat([PSortedRes,pbsres])
    pdosematch=pdosematch.tolist()
    PSortedDoses=PSortedDoses+pdosematch
    sdosematch=ShadowDoses[ShadowDoses==dose]
    sresmatch=Shadow4Res[ShadowDoses==dose]
    sscorematch=ShadowScores[ShadowDoses==dose].tolist()
    SSortedScored=SSortedScored+sscorematch
    sbsres=np.random.choice(sresmatch,(len(sresmatch),1000))
    sbsres=pd.DataFrame(data=sbsres,columns=ColumnNames)
    SSortedRes=pd.concat([SSortedRes,sbsres])
    sdosematch=sdosematch.tolist()
    SSortedDoses=SSortedDoses+sdosematch
    fdosematch=FullDoses[FullDoses==dose]
    fresmatch=Full7Res[FullDoses==dose]
    fscorematch=FullScores[FullDoses==dose].tolist()
    FSortedScored=FSortedScored+fscorematch
    fbsres=np.random.choice(fresmatch,(len(fresmatch),1000))
    fbsres=pd.DataFrame(data=fbsres,columns=ColumnNames)
    FSortedRes=pd.concat([FSortedRes,fbsres])
    fdosematch=fdosematch.tolist()
    FSortedDoses=FSortedDoses+fdosematch
    
#Save the particular bootstrapping instance from May 27, 2020, used to generate the final figure version
PSortedRes.to_csv('FINAL VERSION Bootstrapped Proximal Data.csv')
FSortedRes.to_csv('FINAL VERSION Bootstrapped Full Length Data.csv')
SSortedRes.to_csv('FINAL VERSION Bootstrapped Distal Data.csv')

#Use the bootstrapped residuals to fit curves, along with initial guesses for parameter values
ProxPO=[0.41,-4.9,-0.30,0.040]
ShadowPO=[0.34,-1.5,-1.2,0.036]
FullPO=[1.0,-3.5,-1.0,0.58,-5.2,-0.22,0.061]

PSortedParamAs=[]
PSortedParamBs=[]
PSortedParamCs=[]
PSortedParamDs=[]
PFailed=pd.DataFrame(data=PSortedDoses,columns=['LogDose'])
SSortedParamAs=[]
SSortedParamBs=[]
SSortedParamCs=[]
SSortedParamDs=[]
SFailed=pd.DataFrame(data=SSortedDoses,columns=['LogDose'])
FSortedParamAs=[]
FSortedParamBs=[]
FSortedParamCs=[]
FSortedParamDs=[]
FSortedParamEs=[]
FSortedParamFs=[]
FSortedParamGs=[]
FFailed=pd.DataFrame(data=FSortedDoses,columns=['LogDose'])
for iteration in range(1000):
    pres=PSortedRes.iloc[:,iteration].tolist()
    pbsdata=np.add(pres,PSortedScored)
    try:
        pbsopt,pbscov=curve_fit(logistic4,PSortedDoses,pbsdata,p0=ProxPO,bounds=Bounds4)
        PSortedParamAs.append(pbsopt[0])
        PSortedParamBs.append(pbsopt[1])
        PSortedParamCs.append(pbsopt[2])
        PSortedParamDs.append(pbsopt[3])
    except:
        PFailed['Iter'+str(iteration+1)]=pbsdata
    sres=SSortedRes.iloc[:,iteration].tolist()
    sbsdata=np.add(sres,SSortedScored)
    try:
        sbsopt,sbscov=curve_fit(logistic4,SSortedDoses,sbsdata,p0=ShadowPO,bounds=Bounds4)
        SSortedParamAs.append(sbsopt[0])
        SSortedParamBs.append(sbsopt[1])
        SSortedParamCs.append(sbsopt[2])
        SSortedParamDs.append(sbsopt[3])
    except:
        SFailed['Iter'+str(iteration+1)]=sbsdata
    fres=FSortedRes.iloc[:,iteration].tolist()
    fbsdata=np.add(fres,FSortedScored)
    try:
        fbsopt,fbscov=curve_fit(logistic7,FSortedDoses,fbsdata,p0=FullPO,bounds=Bounds7)
        FSortedParamAs.append(fbsopt[0])
        FSortedParamBs.append(fbsopt[1])
        FSortedParamCs.append(fbsopt[2])
        FSortedParamDs.append(fbsopt[3])
        FSortedParamEs.append(fbsopt[4])
        FSortedParamFs.append(fbsopt[5])
        FSortedParamGs.append(fbsopt[6])
    except:
        FFailed['Iter'+str(iteration+1)]=fbsdata

#Make and save plots of the distributions of bootstrapped parameters
PSortedParams=[PSortedParamAs,PSortedParamBs,PSortedParamCs,PSortedParamDs]
SSortedParams=[SSortedParamAs,SSortedParamBs,SSortedParamCs,SSortedParamDs]
FSortedParams=[FSortedParamAs,FSortedParamBs,FSortedParamCs,FSortedParamDs,FSortedParamEs,FSortedParamFs,FSortedParamGs]
MonoNames=['Top Plateau','Hill Coefficient','EC50','Bottom Plateau']
BiNames=['Top Plateau','Hill Coefficient A','EC50 A','Middle Plateau','Hill Coefficient B','EC50 B','Bottom Plateau']
StartBins=np.linspace(0.2,1.9,num=30)
HCBins=np.linspace(-50,0,num=30)
ECBins=np.linspace(-2.2,0,num=30)
StopBins=np.linspace(0,0.09,num=30)
PlateauBins=np.linspace(0.3,0.9,num=30)
MonoBinList=[StartBins,HCBins,ECBins,StopBins]
BiBinList=[StartBins,HCBins,ECBins,PlateauBins,HCBins,ECBins,StopBins]

for param,name,bins in zip(PSortedParams,MonoNames,MonoBinList):
    fig,ax=plt.subplots(figsize=(8,6))
    fig=plt.hist(param,bins=bins,color='b',density=True)
    plt.setp(ax,ylabel='Bin Density',xlabel='Parameter Value',title='Proximal '+name)
    plt.savefig('FINAL VERSION Proximal Bootstrapped Distributions of '+name+'.pdf',transparent=True)

for param,name,bins in zip(SSortedParams,MonoNames,MonoBinList):
    fig,ax=plt.subplots(figsize=(8,6))
    fig=plt.hist(param,bins=bins,color='m',density=True)
    plt.setp(ax,ylabel='Bin Density',xlabel='Parameter Value',title='Distal '+name)
    plt.savefig('FINAL VERSION Distal Bootstrapped Distributions of '+name+'.pdf',transparent=True)

for param,name,bins in zip(FSortedParams,BiNames,BiBinList):
    fig,ax=plt.subplots(figsize=(8,6))
    fig=plt.hist(param,bins=bins,color='g',density=True)
    plt.setp(ax,ylabel='Bin Density',xlabel='Parameter Value',title='Full Length '+name)
    plt.savefig('FINAL VERSION Full Length Bootstrapped Distributions of '+name+'.pdf',transparent=True)

#Save a table containing the mean and 95% CI bounds for each parameter from the bootstrapped residual population
ReporterParamList=['Proximal','Proximal','Proximal','Proximal','Distal','Distal','Distal','Distal','Full Length','Full Length','Full Length','Full Length','Full Length','Full Length','Full Length']
ParamList=MonoNames+MonoNames+BiNames
ParamPopulations=PSortedParams+SSortedParams+FSortedParams
CILowerList=[]
MedianList=[]
CIUpperList=[]
MeanList=[]
for population in ParamPopulations:
    cilower=np.percentile(population,2.5)
    median=np.median(population)
    ciupper=np.percentile(population,97.5)
    mean=np.mean(population)
    CILowerList.append(cilower)
    MedianList.append(median)
    CIUpperList.append(ciupper)
    MeanList.append(mean)
ParamTable=pd.DataFrame()
ParamTable['Reporter']=ReporterParamList
ParamTable['Parameter']=ParamList
ParamTable['95% CI Median Lower Limit']=CILowerList
ParamTable['Median']=MedianList
ParamTable['95% CI Median Upper Limit']=CIUpperList
ParamTable['Mean']=MeanList
ParamTable.to_csv('FINAL VERSION Parameter Estimate Table.csv')

#Make and save a table of parameter estimates and 95% CIs based on the standard curve fitting
NaiveReporterList=['Proximal','Proximal','Proximal','Proximal','Proximal','Proximal','Proximal','Proximal','Proximal','Proximal','Proximal','Proximal','Proximal','Distal','Distal','Distal','Distal','Distal','Distal','Distal','Distal','Distal','Distal','Distal','Distal','Distal','Full Length','Full Length','Full Length','Full Length','Full Length','Full Length','Full Length','Full Length','Full Length','Full Length','Full Length','Full Length','Full Length']
NaiveTypeList=['Linear','Linear','Monophasic','Monophasic','Monophasic','Monophasic','Biphasic','Biphasic','Biphasic','Biphasic','Biphasic','Biphasic','Biphasic','Linear','Linear','Monophasic','Monophasic','Monophasic','Monophasic','Biphasic','Biphasic','Biphasic','Biphasic','Biphasic','Biphasic','Biphasic','Linear','Linear','Monophasic','Monophasic','Monophasic','Monophasic','Biphasic','Biphasic','Biphasic','Biphasic','Biphasic','Biphasic','Biphasic']
NaiveNameList=['Slope','Intercept','Top Plateau','Hill Coefficient','EC50','Bottom Plateau','Top Plateau','Hill Coefficient A','EC50 A','Middle Plateau','Hill Coefficient B','EC50 B','Bottom Plateau','Slope','Intercept','Top Plateau','Hill Coefficient','EC50','Bottom Plateau','Top Plateau','Hill Coefficient A','EC50 A','Middle Plateau','Hill Coefficient B','EC50 B','Bottom Plateau','Slope','Intercept','Top Plateau','Hill Coefficient','EC50','Bottom Plateau','Top Plateau','Hill Coefficient A','EC50 A','Middle Plateau','Hill Coefficient B','EC50 B','Bottom Plateau']
NaiveEstList=[ProxLinOpt[0],ProxLinOpt[1],Prox4Opt[0],Prox4Opt[1],Prox4Opt[2],Prox4Opt[3],Prox7Opt[0],Prox7Opt[1],Prox7Opt[2],Prox7Opt[3],Prox7Opt[4],Prox7Opt[5],Prox7Opt[6],ShadowLinOpt[0],ShadowLinOpt[1],Shadow4Opt[0],Shadow4Opt[1],Shadow4Opt[2],Shadow4Opt[3],Shadow7Opt[0],Shadow7Opt[1],Shadow7Opt[2],Shadow7Opt[3],Shadow7Opt[4],Shadow7Opt[5],Shadow7Opt[6],FullLinOpt[0],FullLinOpt[1],Full4Opt[0],Full4Opt[1],Full4Opt[2],Full4Opt[3],Full7Opt[0],Full7Opt[1],Full7Opt[2],Full7Opt[3],Full7Opt[4],Full7Opt[5],Full7Opt[6]]
NaiveErrList=[ProxLinErr[0],ProxLinErr[1],Prox4Err[0],Prox4Err[1],Prox4Err[2],Prox4Err[3],Prox7Err[0],Prox7Err[1],Prox7Err[2],Prox7Err[3],Prox7Err[4],Prox7Err[5],Prox7Err[6],ShadowLinErr[0],ShadowLinErr[1],Shadow4Err[0],Shadow4Err[1],Shadow4Err[2],Shadow4Err[3],Shadow7Err[0],Shadow7Err[1],Shadow7Err[2],Shadow7Err[3],Shadow7Err[4],Shadow7Err[5],Shadow7Err[6],FullLinErr[0],FullLinErr[1],Full4Err[0],Full4Err[1],Full4Err[2],Full4Err[3],Full7Err[0],Full7Err[1],Full7Err[2],Full7Err[3],Full7Err[4],Full7Err[5],Full7Err[6]]
NaiveTable=pd.DataFrame()
NaiveTable['Reporter']=NaiveReporterList
NaiveTable['Curve Type']=NaiveTypeList
NaiveTable['Parameter']=NaiveNameList
NaiveTable['Parameter Estimate']=NaiveEstList
NaiveTable['Standard Error']=NaiveErrList
NaiveTable.to_csv('FINAL VERSION Naive Parameter Estimate Table.csv')

#Plot out families of curves for each reporter
#Split param table by reporter, and isolate sets of parameter options
ProxParams=ParamTable[ParamTable['Reporter']=='Proximal']
DistalParams=ParamTable[ParamTable['Reporter']=='Distal']
FullParams=ParamTable[ParamTable['Reporter']=='Full Length']

ProxStart=ProxParams.iloc[0,2:]
ProxHC=ProxParams.iloc[1,2:]
ProxEC=ProxParams.iloc[2,2:]
ProxStop=ProxParams.iloc[3,2:]
DistalStart=DistalParams.iloc[0,2:]
DistalHC=DistalParams.iloc[1,2:]
DistalEC=DistalParams.iloc[2,2:]
DistalStop=DistalParams.iloc[3,2:]
FullStart=FullParams.iloc[0,2:]
FullHCA=FullParams.iloc[1,2:]
FullECA=FullParams.iloc[2,2:]
FullPlateau=FullParams.iloc[3,2:]
FullHCB=FullParams.iloc[4,2:]
FullECB=FullParams.iloc[5,2:]
FullStop=FullParams.iloc[6,2:]

#Set up lines used in legends of the plots below
fitline=mlines.Line2D([],[],color='gray',label='Bootstrap Parameter Fits',ls=':')
pfitline=mlines.Line2D([],[],color='b',label='Proximal Parameter Fits',ls=':')
dfitline=mlines.Line2D([],[],color='m',label='Distal Parameter Fits',ls=':')
ffitline=mlines.Line2D([],[],color='g',label='Full Length Parameter Fits',ls=':')

#Plot for just the proximal family fits
fig,ax=plt.subplots(figsize=(8,6))
for start in ProxStart:
    for hc in ProxHC:
        for ec in ProxEC:
            for stop in ProxStop:
                fig=plt.plot(detailedx,peval4(detailedx,[start,hc,ec,stop]),'gray',ls=':')
fig=sns.lineplot(x='ModLogDose',y="NSSEVNormedToFL0.01581",color="b",ax=ax,ci=95,data=Prox)
fig=plt.xlabel("Log[U0126 (uM)]")
fig=plt.ylabel("Normalized Expression")
fig=plt.title('Proximal Bootstrap Parameter Fits')
fig=plt.legend(handles=[pline,fitline])
fig=plt.savefig('FINAL VERSION Proximal Bootstrap Parameter Fit Family.pdf',transparent=True)

#Plot for just the distal family fits
fig,ax=plt.subplots(figsize=(8,6))
for start in DistalStart:
    for hc in DistalHC:
        for ec in DistalEC:
            for stop in DistalStop:
                fig=plt.plot(detailedx,peval4(detailedx,[start,hc,ec,stop]),'gray',ls=':')
fig=sns.lineplot(x='ModLogDose',y="NSSEVNormedToFL0.01581",color="m",ax=ax,ci=95,data=Shadow)
fig=plt.xlabel("Log[U0126 (uM)]")
fig=plt.ylabel("Normalized Expression")
fig=plt.title('Distal Bootstrap Parameter Fits')
fig=plt.legend(handles=[dline,fitline])
fig=plt.savefig('FINAL VERSION Distal Bootstrap Parameter Fit Family.pdf',transparent=True)

#Plot comparing the proximal and distal family fits
fig,ax=plt.subplots(figsize=(8,6))
for pstart,dstart in zip(ProxStart,DistalStart):
    for phc,dhc in zip(ProxHC,DistalHC):
        for pec,dec in zip(ProxEC,DistalEC):
            for pstop,dstop in zip(ProxStop,DistalStop):
                fig=plt.plot(detailedx,peval4(detailedx,[pstart,phc,pec,pstop]),'b',ls=':')
                fig=plt.plot(detailedx,peval4(detailedx,[dstart,dhc,dec,dstop]),'m',ls=':')
fig=plt.xlabel("Log[U0126 (uM)]")
fig=plt.ylabel("Normalized Expression")
fig=plt.title('Proximal and Distal Bootstrap Parameter Fits Comparison')
fig=plt.legend(handles=[pfitline,dfitline])
fig=plt.savefig('FINAL VERSION Proximal and Distal Parameter Fit Family Comparison.pdf',transparent=True)

#Plot for just the full length family fits
fig,ax=plt.subplots(figsize=(8,6))
for start in FullStart:
    for hca in FullHCA:
        for eca in FullECA:
            for plateau in FullPlateau:
                for hcb in FullHCB:
                    for ecb in FullECB:
                        for stop in FullStop:
                            fig=plt.plot(detailedx,peval7(detailedx,[start,hca,eca,plateau,hcb,ecb,stop]),'gray',ls=':')
fig=sns.lineplot(x='ModLogDose',y="NSSEVNormedToFL0.01581",color="g",ax=ax,ci=95,data=Full)
fig=plt.xlabel("Log[U0126 (uM)]")
fig=plt.ylabel("Normalized Expression")
fig=plt.title('Full Length Bootstrap Parameter Fits')
plt.setp(ax,ylim=(-0.02,1.6))
fig=plt.legend(handles=[fline,fitline])
fig=plt.savefig('FINAL VERSION Full Length Bootstrap Parameter Fit Family.pdf',transparent=True)

#Plot all three families together
fig,ax=plt.subplots(figsize=(8,6))
for pstart,dstart in zip(ProxStart,DistalStart):
    for phc,dhc in zip(ProxHC,DistalHC):
        for pec,dec in zip(ProxEC,DistalEC):
            for pstop,dstop in zip(ProxStop,DistalStop):
                fig=plt.plot(detailedx,peval4(detailedx,[pstart,phc,pec,pstop]),'b',ls=':')
                fig=plt.plot(detailedx,peval4(detailedx,[dstart,dhc,dec,dstop]),'m',ls=':')
for start in FullStart:
    for hca in FullHCA:
        for eca in FullECA:
            for plateau in FullPlateau:
                for hcb in FullHCB:
                    for ecb in FullECB:
                        for stop in FullStop:
                            fig=plt.plot(detailedx,peval7(detailedx,[start,hca,eca,plateau,hcb,ecb,stop]),'g',ls=':')
fig=plt.xlabel("Log[U0126 (uM)]")
fig=plt.ylabel("Normalized Expression")
fig=plt.title('Bootstrap Parameter Fits Comparison')
plt.setp(ax,ylim=(-0.02,1.6))
fig=plt.legend(handles=[pfitline,dfitline,ffitline])
fig=plt.savefig('FINAL VERSION All Reporters Parameter Fit Family Comparison.pdf',transparent=True)

#Plot single best fit curves from bootstrapped values (using median value) for prox and shadow over the SEV data
ProxBest=ProxParams.Median.tolist()
DistalBest=DistalParams.Median.tolist()
PDScores=SEVScores[SEVScores['enhancer_construct']!='Shadow+Spacer+Proximal']
pline=mlines.Line2D([],[],color='b',label='Proximal')

fig,ax=plt.subplots(figsize=(8,6))
fig=sns.lineplot(x="ModLogDose",y="NSSEVNormedToFL0.01581",hue="enhancer_construct",palette=["blue","magenta"],markers=['o','o'],dashes=False,style='enhancer_construct',ax=ax,ci=95,data=PDScores)
fig=plt.plot(detailedx,peval4(detailedx,ProxBest),':b')
fig=plt.plot(detailedx,peval4(detailedx,DistalBest),':m')
fig=plt.plot(ProxBest[2],peval4(ProxBest[2],ProxBest),'ko')
fig=plt.plot(DistalBest[2],peval4(DistalBest[2],DistalBest),'ko')
fig=plt.hlines(peval4(ProxBest[2],ProxBest),ProxEC[0],ProxEC[2],'k','-')
fig=plt.hlines(peval4(DistalBest[2],DistalBest),DistalEC[0],DistalEC[2],'k','-')
fig=plt.xticks(ticks=doseticks,labels=doselabels)
fig=plt.xlabel("[U0126 (uM)]")
fig=plt.ylabel("Normalized Expression")
fig=plt.title('SEV data and best curve fits')
plt.setp(ax,ylim=(-0.02,0.7))
fig=plt.legend(handles=[pline,dline,pfitline,dfitline])
fig=plt.savefig('FINAL VERSION SEV Data and Curve Fit for Prox and Distal Plot.pdf',transparent=True)

#Plot FL bootstrapped median curves, making two curves in comparison with one another.
#Both will use median values at every parameter, except the HCs will be plotted separately.  Cutoffs are -15 for HCA and -12 for HCB
FullBest=FullParams.Median.tolist()
FHCBHigh=[]
FHCBLow=[]
FHCEHigh=[]
FHCELow=[]
for param in FSortedParamBs:
    if param<-15:
        FHCBHigh.append(param)
    else:
        FHCBLow.append(param)
for param in FSortedParamEs:
    if param<-12:
        FHCEHigh.append(param)
    else:
        FHCELow.append(param)
FHCBHighL,FHCBHighU=np.percentile(FHCBHigh,[2.5,97.5])
FHCBLowL,FHCBLowU=np.percentile(FHCBLow,[2.5,97.5])
FHCEHighL,FHCEHighU=np.percentile(FHCEHigh,[2.5,97.5])
FHCELowL,FHCELowU=np.percentile(FHCELow,[2.5,97.5])
FHCBHighMean=np.average(FHCBHigh)
FHCBLowMean=np.average(FHCBLow)
FHCEHighMean=np.average(FHCEHigh)
FHCELowMean=np.average(FHCELow)
FHCBHighMed=np.median(FHCBHigh)
FHCBLowMed=np.median(FHCBLow)
FHCEHighMed=np.median(FHCEHigh)
FHCELowMed=np.median(FHCELow)
FullHillCompare=pd.DataFrame()
FullHillCompare['Coefficient']=['HCA Steep','HCA Shallow','HCB Steep','HCB Shallow']
FullHillCompare['Median']=[FHCBHighMed,FHCBLowMed,FHCEHighMed,FHCELowMed]
FullHillCompare['95% CI Lower']=[FHCBHighL,FHCBLowL,FHCEHighL,FHCELowL]
FullHillCompare['95% CI Upper']=[FHCBHighU,FHCBLowU,FHCEHighU,FHCELowU]
FullHillCompare['Mean']=[FHCBHighMean,FHCBLowMean,FHCEHighMean,FHCELowMean]
FullHillCompare.to_csv('FINAL VERSION FL Hill Coefficient Table.csv')

steep=mlines.Line2D([],[],color='g',ls=':',label='Steep Solution')
likely=mlines.Line2D([],[],color='g',ls='--',label='Shallow Solution')

fig,ax=plt.subplots(figsize=(8,6))
fig=sns.lineplot(x="ModLogDose",y="NSSEVNormedToFL0.01581",color='g',markers=['o'],dashes=False,style='enhancer_construct',ax=ax,ci=95,data=Full)
fig=plt.plot(detailedx,peval7(detailedx,[FullBest[0],FHCBHighMed,FullBest[2],FullBest[3],FHCEHighMed,FullBest[5],FullBest[6]]),':g')
fig=plt.plot(detailedx,peval7(detailedx,[FullBest[0],FHCBLowMed,FullBest[2],FullBest[3],FHCELowMed,FullBest[5],FullBest[6]]),'--g')
fig=plt.plot(FullBest[2],peval7(FullBest[2],FullBest),'ko')
fig=plt.plot(FullBest[5],peval7(FullBest[5],FullBest),'ko')
fig=plt.hlines(peval7(FullBest[2],FullBest),FullECA[0],FullECA[2],'k','-')
fig=plt.hlines(peval7(FullBest[5],FullBest),FullECB[0],FullECB[2],'k','-')
fig=plt.xticks(ticks=doseticks,labels=doselabels)
fig=plt.xlabel('[U0126 (uM)]')
fig=plt.ylabel('Normalized Expression')
fig=plt.title('FL Hill Coefficient Comparison')
plt.setp(ax,ylim=(-0.02,1.5))
fig=plt.legend(handles=[fline,steep,likely])
fig=plt.savefig('FINAL VERSION FL Hill Coefficient Comparison.pdf',transparent=True)

#Calculate expanded AIC and RL values on the FL curves, including the naive and the bootstrapped curves
SteepRes=residuals7([FullBest[0],FHCBHighMed,FullBest[2],FullBest[3],FHCEHighMed,FullBest[5],FullBest[6]],FullScores,FullDoses)
ShallowRes=residuals7([FullBest[0],FHCBLowMed,FullBest[2],FullBest[3],FHCELowMed,FullBest[5],FullBest[6]],FullScores,FullDoses)

SteepRSS=np.sum(SteepRes*SteepRes)
ShallowRSS=np.sum(ShallowRes*ShallowRes)

SteepAIC=FullN*np.log(SteepRSS/FullN)+(2*8)
ShallowAIC=FullN*np.log(ShallowRSS/FullN)+(2*8)

MinFullAIC=min(FullLinAIC,Full4AIC,Full7AIC,SteepAIC,ShallowAIC)

NewFullLinRL=math.exp((MinFullAIC-FullLinAIC)/2)
NewFull4RL=math.exp((MinFullAIC-Full4AIC)/2)
NewFull7RL=math.exp((MinFullAIC-Full7AIC)/2)
SteepRL=math.exp((MinFullAIC-SteepAIC)/2)
ShallowRL=math.exp((MinFullAIC-ShallowAIC)/2)

RLList=[ProxLinRL,Prox4RL,Prox7RL,ShadowLinRL,Shadow4RL,Shadow7RL,FullLinRL,Full4RL,Full7RL]
FLCurveSelection=pd.DataFrame(columns=['CurveType','AIC','RelativeLikelihood'])
FLCurveSelection.iloc[:,0]=['Linear','Monophasic','Naive Biphasic','Steep Biphasic','Shallow Biphasic']
FLCurveSelection.iloc[:,1]=[FullLinAIC,Full4AIC,Full7AIC,SteepAIC,ShallowAIC]
FLCurveSelection.iloc[:,2]=[NewFullLinRL,NewFull4RL,NewFull7RL,SteepRL,ShallowRL]
FLCurveSelection.to_csv("FINAL VERSION FL Curve Selection Table.csv")

#Plot single best fit curves from bootstrapped values (median parameter) for all reporters over the SEV data
fig,ax=plt.subplots(figsize=(8,6))
fig=sns.lineplot(x="ModLogDose",y="NSSEVNormedToFL0.01581",hue="enhancer_construct",palette=["blue","green","magenta"],ax=ax,ci=95,data=SEVScores)
fig=plt.plot(detailedx,peval4(detailedx,ProxBest),':b')
fig=plt.plot(detailedx,peval4(detailedx,DistalBest),':m')
fig=plt.plot(detailedx,peval7(detailedx,FullBest),':g')
fig=plt.xlabel("Log[U0126 (uM)]")
fig=plt.ylabel("Normalized Expression")
fig=plt.title('SEV data and best curve fits')
plt.setp(ax,ylim=(-0.02,1.45))
fig=plt.legend(handles=[pline,dline,fline,pfitline,dfitline,ffitline])
fig=plt.savefig('FINAL VERSION SEV Data and Curve Fit for all Reporters Plot.pdf',transparent=True)

#Plot a modified version of the curves for all reporters, but this time sub the 95% CI upper limit for just the HCs on full length to make it more biologically likely
FullBestMod=FullBest
FullBestMod[1]=CIUpperList[9]
FullBestMod[4]=CIUpperList[12]

fig,ax=plt.subplots(figsize=(8,6))
fig=sns.lineplot(x="ModLogDose",y="NSSEVNormedToFL0.01581",hue="enhancer_construct",palette=["blue","green","magenta"],ax=ax,ci=95,data=SEVScores)
fig=plt.plot(detailedx,peval4(detailedx,ProxBest),':b')
fig=plt.plot(detailedx,peval4(detailedx,DistalBest),':m')
fig=plt.plot(detailedx,peval7(detailedx,FullBestMod),':g')
fig=plt.xlabel("Log[U0126 (uM)]")
fig=plt.ylabel("Normalized Expression")
fig=plt.title('SEV data and best curve fits')
fig=plt.legend(handles=[pline,dline,fline,pfitline,dfitline,ffitline])
plt.setp(ax,ylim=(-0.02,1.45))
fig=plt.savefig('FINAL VERSION SEV Data and Curve Fit for all Reporters Plot with Modded Full Length.pdf',transparent=True)

#Run t-tests on the proximal vs distal enhancer parameter distributions and obtain CI of the difference in parameters
TStatList=[]
PValList=[]
for pparam,dparam in zip(PSortedParams,SSortedParams):
    tstat,pval=ttest_ind(pparam,dparam)
    TStatList.append(tstat)
    PValList.append(pval)
PDComparison=pd.DataFrame()
PDComparison['Parameter']=MonoNames
PDComparison['T-Statistic']=TStatList
PDComparison['p-value']=PValList
PDComparison.to_csv('FINAL VERSION t-test results for Proximal and Distal Comparison.csv')

#Run Wilcoxon rank-sum tests on the same distributions, which do not assume Gaussian distributions for data
WilcoxonList=[]
WilcoxonPValList=[]
for pparam,dparam in zip(PSortedParams,SSortedParams):
    tstat,pval=ranksums(pparam,dparam)
    WilcoxonList.append(tstat)
    WilcoxonPValList.append(pval)
PDWilcoxonComparison=pd.DataFrame()
PDWilcoxonComparison['Parameter']=MonoNames
PDWilcoxonComparison['Test Statistic']=WilcoxonList
PDWilcoxonComparison['p-value']=WilcoxonPValList
PDWilcoxonComparison.to_csv('FINAL VERSION Wilcoxon Test results for Proximal and Distal Comparison.csv')

#Run t-tests to compare first phase full length parameters to distal, and second phase full length parameters to proximal
#For the distal/phase one comparison, phase one A and D need to be recalculated to be testing the magnitude of the difference between them
#Because the magnitude is used for distal, also run an equivalent test on proximal
FTStatList=[]
FPValList=[]
FComparisonList=['Phase One/Distal Hill Coefficient','Phase One/Distal EC50','Phase One/Distal Magnitude','Phase Two/Proximal Start','Phase Two/Proximal Hill Coefficient','Phase Two/Proximal EC50','Phase Two/Proximal Stop','Phase Two/Proximal Magnitude']
FOneChanges=[]
for a,d in zip(FSortedParamAs,FSortedParamDs):
    change=a-d
    FOneChanges.append(change)
FTwoChanges=[]
for d,g in zip(FSortedParamDs,FSortedParamGs):
    change=d-g
    FTwoChanges.append(change)
FLPhaseOneParams=[FSortedParamBs,FSortedParamCs,FOneChanges]
FLPhaseTwoParams=[FSortedParamDs,FSortedParamEs,FSortedParamFs,FSortedParamGs,FTwoChanges]
DChanges=[]
for a,d in zip(SSortedParamAs,SSortedParamDs):
    change=a-d
    DChanges.append(change)
DCompParams=[SSortedParamBs,SSortedParamCs,DChanges]
PChanges=[]
for a,d in zip(PSortedParamAs,PSortedParamDs):
    change=a-d
    PChanges.append(change)
PCompParams=[PSortedParamAs,PSortedParamBs,PSortedParamCs,PSortedParamDs,PChanges]
for oparam,dparam in zip(FLPhaseOneParams,DCompParams):
    tstat,pval=ttest_ind(oparam,dparam)
    FTStatList.append(tstat)
    FPValList.append(pval)
for tparam,pparam in zip(FLPhaseTwoParams,PCompParams):
    tstat,pval=ttest_ind(tparam,pparam)
    FTStatList.append(tstat)
    FPValList.append(pval)
FComparison=pd.DataFrame()
FComparison['Parameters']=FComparisonList
FComparison['T-Statistic']=FTStatList
FComparison['p-value']=FPValList
FComparison.to_csv('FINAL VERSION t-test results for Comparison to FL.csv')

#Also run Wilcoxon rank-sum tests on the same set of full length/individual enhancer comparisons
FWilcoxList=[]
FWilcoxPvalList=[]
for oparam,dparam in zip(FLPhaseOneParams,DCompParams):
    tstat,pval=ranksums(oparam,dparam)
    FWilcoxList.append(tstat)
    FWilcoxPvalList.append(pval)
for tparam,pparam in zip(FLPhaseTwoParams,PCompParams):
    tstat,pval=ranksums(tparam,pparam)
    FWilcoxList.append(tstat)
    FWilcoxPvalList.append(pval)
FWilcox=pd.DataFrame()
FWilcox['Parameters']=FComparisonList
FWilcox['Test Statistic']=FWilcoxList
FWilcox['p-value']=FWilcoxPvalList
FWilcox.to_csv('FINAL VERSION Wilcoxon Test results for Comparison to FL.csv')
```
