## Supplemental Appendix S3 for "*Ciona Brachyury* proximal and distal enhancers have different FGF dose-response relationships": Fig 3 and Associated Supplement.py

```
#!/usr/bin/env python3
# -*- coding: utf-8 -*-
"""
Created on Thu May 28 16:48:02 2020

@author: Matt
"""

#Built to be the final repository of code for all things to do with the Cell Measurement plotting
#This is NOT a place to tinker with ideas or generate completely new plots
#ONLY use this to make adjustments for final versions of figure panels, and as a script to produce them

#Import useful libraries and settings
import numpy as np
import pandas as pd
import seaborn as sns
import matplotlib.pyplot as plt
import os
import matplotlib.lines as mlines
from scipy.stats import f_oneway
from scipy.stats import ttest_1samp
from statsmodels.stats.multicomp import MultiComparison
plt.rcParams['pdf.fonttype']=42

#Read in SEV and cell measure data, add a serial number to cell measure data
os.chdir("/Volumes/MH Veeman Lab Home Backup/Cell Measurements data and output")
Measurements=pd.read_csv("Cell Measurements MH edits.csv")
Serial=[]
Num=1
for cell in range(len(Measurements)):
    Serial.append(Num)
    Num=Num+1
Measurements['SerialNum']=Serial

#Use mode as a way to select on/off status in cell measurements.
#The use of mode and the particular threshold were selected based on graphs from CMII
OnOff=[]
for cell in range(len(Measurements)):
    if Measurements.Mode[cell]<10.5:
        OnOff.append(0)
    else:
        OnOff.append(1)
Measurements['OnOff']=OnOff

#Experiment normalize and scale each measure in the same manner as done for SEV
ReporterList=['Prox','Distal','FL']
XPList=[1,2,3,4,5,6,11,12,13,14,15,16,17,18]
RawMeasureList=['Mean','StDev','Mode','Min','Max','IntDen','Median','RawIntDen']
Reps=[]
XPS=[]
Factors=[[],[],[],[],[],[],[],[]]
NormalizationAverages=pd.DataFrame()
for reporter in ReporterList:
    data=Measurements[Measurements['Reporter']==reporter]
    for xp in XPList:
        xpdata=data[data['Experiment Number']==xp]
        dosedata=xpdata[xpdata['Dose']=='DMSO']
        Reps.append(reporter)
        XPS.append(xp)
        for measure,location in zip(RawMeasureList,range(len(Factors))):
            measuredata=np.average(dosedata[measure])
            Factors[location].append(measuredata)
NormalizationAverages['Reporter']=Reps
NormalizationAverages['XP']=XPS
for measurement,name in zip(Factors,RawMeasureList):
    NormalizationAverages[name]=measurement
    
NSfactors=[[],[],[],[],[],[],[],[]]
NSAverages=pd.DataFrame()
for reporter in ReporterList:
    data=Measurements[Measurements['Reporter']==reporter]
    scaledata=data[data['Dose']=='DMSO']
    for measure,location in zip(RawMeasureList,range(len(NSfactors))):
        measuredata=np.average(scaledata[measure])
        NSfactors[location].append(measuredata)
NSAverages['Reporter']=ReporterList
for measurement,name in zip(NSfactors,RawMeasureList):
    NSAverages[name]=measurement
    
NMeasures=[[],[],[],[],[],[],[],[]]
NSMeasures=[[],[],[],[],[],[],[],[]]
NNSMeasures=[[],[],[],[],[],[],[],[]]
for cell in range(len(Measurements)):
    xp=Measurements.iloc[cell,0]
    reporter=Measurements.Reporter[cell]
    control=NormalizationAverages[NormalizationAverages['Reporter']==reporter]
    control=control[control['XP']==xp]
    for measure,position in zip(RawMeasureList,range(len(RawMeasureList))):
        normal=np.average(control[measure])
        data=Measurements[measure]
        raw=data[cell]
        Nmeasure=raw/normal
        NMeasures[position].append(Nmeasure)
        scale=NSAverages[NSAverages['Reporter']==reporter]
        scale=np.average(scale[measure])
        NSMeasure=Nmeasure*scale
        NSMeasures[position].append(NSMeasure)
        norm=NSAverages[measure]
        norm=norm[2]
        NNSMeasure=NSMeasure/norm
        NNSMeasures[position].append(NNSMeasure)

for one,two,three,name,position in zip(NMeasures,NSMeasures,NNSMeasures,RawMeasureList,range(len(RawMeasureList))):
    Measurements['XPNormed'+name]=NMeasures[position]
    Measurements['XPNormedScaled'+name]=NSMeasures[position]
    Measurements['FLNormed'+name]=NNSMeasures[position]

#Useful set of lists to be referenced multiple times over the course of the plotting:
LongReporterList=['Prox','Distal','FL','Prox','Distal','FL','Prox','Distal','FL']
DoseList=['DMSO','0.1','0.342']
LongDoseList=['DMSO','DMSO','DMSO','0.1','0.1','0.1','0.342','0.342','0.342']
MeasurementsList=['Mean','StDev','Mode','Min','Max','IntDen','Median','RawIntDen']
NormedMeasurementsList=['XPNormedScaledMean','XPNormedScaledStDev','XPNormedScaledMode','XPNormedScaledMin','XPNormedScaledMax','XPNormedScaledIntDen','XPNormedScaledMedian','XPNormedScaledRawIntDen']

#Generate a scatterplot for mode v normalized mean to show the on/off distinction
LogMode=[]
LogMeans=[]
Modes=Measurements['Mode']
NMeans=Measurements['XPNormedScaledMean']
for cell in range(len(Modes)):
    logmode=np.log2(Modes[cell]+1)
    logmean=np.log2(NMeans[cell]+1)
    LogMode.append(logmode)
    LogMeans.append(logmean)
Measurements['LogMode']=LogMode
Measurements['LogMeans']=LogMeans
fig=plt.figure()
fig=sns.jointplot(x='LogMode',y='LogMeans',data=Measurements)
fig.savefig('first pass mode and mean scatterplot.pdf',transparent=True)
OnCells=Measurements[Measurements['OnOff']==1]

#Set up variables and lists to be used in the generation of the plot for determining on/off status
Prox=Measurements[Measurements['Reporter']=='Prox']
Distal=Measurements[Measurements['Reporter']=='Distal']
Full=Measurements[Measurements['Reporter']=='FL']
ProxControl=Prox[Prox['Dose']=='DMSO']
DistalControl=Distal[Distal['Dose']=='DMSO']
FullControl=Full[Full['Dose']=='DMSO']
ProxPointOne=Prox[Prox['Dose']=='0.1']
DistalPointOne=Distal[Distal['Dose']=='0.1']
FullPointOne=Full[Full['Dose']=='0.1']
ProxPointThree=Prox[Prox['Dose']=='0.342']
DistalPointThree=Distal[Distal['Dose']=='0.342']
FullPointThree=Full[Full['Dose']=='0.342']
ReporterData=[Prox,Distal,Full]

ConditionList=[ProxControl,DistalControl,FullControl,ProxPointOne,DistalPointOne,FullPointOne,ProxPointThree,DistalPointThree,FullPointThree]
RowsAndColumns=[[0,0],[0,1],[0,2],[1,0],[1,1],[1,2],[2,0],[2,1],[2,2]]
meanbins=np.linspace(2,13,num=21)
meanx=meanbins[:-1]
stdevbins=np.linspace(2,11,num=21)
stdevx=stdevbins[:-1]
modebins=np.linspace(1,13,num=21)
modex=modebins[:-1]
minbins=np.linspace(0,13,num=21)
minx=minbins[:-1]
maxbins=np.linspace(4,13,num=21)
maxx=maxbins[:-1]
intdenbins=np.linspace(5,16,num=21)
intdenx=intdenbins[:-1]
medianbins=np.linspace(1,13,num=21)
medianx=medianbins[:-1]
rawintdenbins=np.linspace(8,19,num=21)
rawintdenx=rawintdenbins[:-1]
BinsList=[meanbins,stdevbins,modebins,minbins,maxbins,intdenbins,medianbins,rawintdenbins]
xList=[meanx,stdevx,modex,minx,maxx,intdenx,medianx,rawintdenx]
RawColorList=['-b','-m','-g','-b','-m','-g','-b','-m','-g']
NormColorList=[':b',':m',':g',':b',':m',':g',':b',':m',':g']
yList=[(-0.01,0.45),(-0.01,0.8),(-0.01,0.3),(-0.01,0.2),(-0.01,0.75),(-0.01,0.45),(-0.01,0.3),(-0.01,0.7)]
plineraw=mlines.Line2D([],[],color='b',linestyle='-',label='Prox Raw')
dlineraw=mlines.Line2D([],[],color='m',linestyle='-',label='Distal Raw')
flineraw=mlines.Line2D([],[],color='g',linestyle='-',label='Full Length Raw')
plinenorm=mlines.Line2D([],[],color='b',linestyle=':',label='Prox Norm')
dlinenorm=mlines.Line2D([],[],color='m',linestyle=':',label='Distal Norm')
flinenorm=mlines.Line2D([],[],color='g',linestyle=':',label='Full Length Norm')

#Produce plots of histogram distributions to show selection of mode as the on/off identifier
for raw,norm,binstyle,xstyle,ystyle in zip(MeasurementsList,NormedMeasurementsList,BinsList,xList,yList):
    histlistraw=[]
    histlistnorm=[]
    for condition in ConditionList:
        histogramraw=plt.hist(np.log2(condition[raw]+1),bins=binstyle,density=True)
        histogramnorm=plt.hist(np.log2(condition[norm]+1),bins=binstyle,density=True)
        histlistraw.append(histogramraw[0])
        histlistnorm.append(histogramnorm[0])
    plt.clf()
    fig,ax=plt.subplots(nrows=3,ncols=3,figsize=(16,12))
    for rawhist,normhist,position,rawcolor,normcolor in zip(histlistraw,histlistnorm,RowsAndColumns,RawColorList,NormColorList):
        ax[position[0],position[1]].plot(xstyle,rawhist,rawcolor)
        ax[position[0],position[1]].plot(xstyle,normhist,normcolor)
    plt.setp(ax[0,0],title='Proximal')
    plt.setp(ax[0,1],title='Shadow')
    plt.setp(ax[0,2],title='Full Length')
    plt.setp(ax[0,0],ylabel='DMSO')
    plt.setp(ax[1,0],ylabel='0.1 uM')
    plt.setp(ax[2,0],ylabel='0.342 uM')
    plt.setp(ax,ylim=ystyle)
    plt.legend(handles=[plineraw,dlineraw,flineraw,plinenorm,dlinenorm,flinenorm],loc=(1.05,1.5))
    fig.text(0.5,0.05,'Log2('+raw+')',ha='center',fontsize=15)
    fig.text(0.07,0.5,'Density',va='center',rotation='vertical',fontsize=15)
    fig.savefig('FINAL VERSION ALL DATA All Cells Distributions of '+raw+' with 20 bins and normalization.pdf',transparent=True)

#Bootstrap means and normed means of ALL CELLS, contain by electroporation
ColumnNames=[]
for x in range(1000):
    iteration=x+1
    name='iter'+str(iteration)
    ColumnNames.append(name)

BSMeans=pd.DataFrame()
BSNormedMeans=pd.DataFrame()
FullBSMeans=np.zeros((1,1000))
FullBSNormMeans=np.zeros((1,1000))
FullXPList=[]
FullRepList=[]
FullDoseList=[]
for xp in XPList:
    data=Measurements[Measurements['Experiment Number']==xp]
    for reporter in ReporterList:
        repdata=data[data['Reporter']==reporter]
        for dose in DoseList:
            dosedata=repdata[repdata['Dose']==dose]
            means=dosedata.Mean.tolist()
            normmeans=dosedata.FLNormedMean.tolist()
            if  len(means)>0:
                bsmeans=np.random.choice(means,(len(means),1000))
                bsnormmeans=np.random.choice(normmeans,(len(normmeans),1000))
                xplist=[]
                replist=[]
                doselist=[]
                for cell in range(len(means)):
                    xplist.append(xp)
                    replist.append(reporter)
                    doselist.append(dose)
                FullXPList=FullXPList+xplist
                FullRepList=FullRepList+replist
                FullDoseList=FullDoseList+doselist
                FullBSMeans=np.concatenate((FullBSMeans,bsmeans),axis=0)
                FullBSNormMeans=np.concatenate((FullBSNormMeans,bsnormmeans),axis=0)
FullBSMeans=FullBSMeans[1:,:]
FullBSNormMeans=FullBSNormMeans[1:,:]
                
BSList=[BSMeans,BSNormedMeans]
BSData=[FullBSMeans,FullBSNormMeans]
for df,bsdata in zip(BSList,BSData):
    df['XP']=FullXPList
    df['Reporter']=FullRepList
    df['Dose']=FullDoseList
    for column,location in zip(ColumnNames,range(1000)):
        df[column]=bsdata[:,location]

#Store the data from the particular bootstrapping instance used to generate final plot
BSMeans.to_csv('FINAL VERSION Cell Measurement Means Bootstrapping.csv')
BSNormedMeans.to_csv('FINAL VERSION Cell Measurement Normalized Means Bootstrapping.csv')

#Bootstrap means and normed means of ON CELLS, contain by electroporation
BSOnMeans=pd.DataFrame()
BSOnNormedMeans=pd.DataFrame()
FullBSOnMeans=np.zeros((1,1000))
FullBSOnNormMeans=np.zeros((1,1000))
FullXPOnList=[]
FullRepOnList=[]
FullDoseOnList=[]
for xp in XPList:
    data=OnCells[OnCells['Experiment Number']==xp]
    for reporter in ReporterList:
        repdata=data[data['Reporter']==reporter]
        for dose in DoseList:
            dosedata=repdata[repdata['Dose']==dose]
            means=dosedata.Mean.tolist()
            normmeans=dosedata.FLNormedMean.tolist()
            if  len(means)>0:
                bsmeans=np.random.choice(means,(len(means),1000))
                bsnormmeans=np.random.choice(normmeans,(len(normmeans),1000))
                xplist=[]
                replist=[]
                doselist=[]
                for cell in range(len(means)):
                    xplist.append(xp)
                    replist.append(reporter)
                    doselist.append(dose)
                FullXPOnList=FullXPOnList+xplist
                FullRepOnList=FullRepOnList+replist
                FullDoseOnList=FullDoseOnList+doselist
                FullBSOnMeans=np.concatenate((FullBSOnMeans,bsmeans),axis=0)
                FullBSOnNormMeans=np.concatenate((FullBSOnNormMeans,bsnormmeans),axis=0)
FullBSOnMeans=FullBSOnMeans[1:,:]
FullBSOnNormMeans=FullBSOnNormMeans[1:,:]
                
BSOnList=[BSOnMeans,BSOnNormedMeans]
BSOnData=[FullBSOnMeans,FullBSOnNormMeans]
for df,bsdata in zip(BSOnList,BSOnData):
    df['XP']=FullXPOnList
    df['Reporter']=FullRepOnList
    df['Dose']=FullDoseOnList
    for column,location in zip(ColumnNames,range(1000)):
        df[column]=bsdata[:,location]

#Store the data from the particular bootstrapping instance used to generate final plot
BSOnMeans.to_csv('FINAL VERSION Cell Measurement ON CELLS Means Bootstrapping.csv')
BSOnNormedMeans.to_csv('FINAL VERSION Cell Measurement ON CELLS Normalized Means Bootstrapping.csv')

#Gather mean and 90th percentile of normedmeans from the bootstrap data of ALL CELLS
MeansOfNormedMeans=[]
PercentileOfNormedMeans=[]
OrderedReporters=[]
OrderedDoses=[]
for reporter in ReporterList:
    data=BSNormedMeans[BSNormedMeans['Reporter']==reporter]
    for dose in DoseList:
        dosedata=data[data['Dose']==dose]
        for column in ColumnNames:
            iteration=dosedata[column]
            meanofmeans=np.average(iteration)
            percentileofmeans=np.percentile(iteration,90)
            OrderedReporters.append(reporter)
            OrderedDoses.append(dose)
            MeansOfNormedMeans.append(meanofmeans)
            PercentileOfNormedMeans.append(percentileofmeans)
BootStrapOfNormedMeans=pd.DataFrame()
BootStrapOfNormedMeans['Reporter']=OrderedReporters
BootStrapOfNormedMeans['Dose']=OrderedDoses
BootStrapOfNormedMeans['Mean']=MeansOfNormedMeans
BootStrapOfNormedMeans['Percentile']=PercentileOfNormedMeans

#Gather mean and 90th percentile of normedmeans from the bootstrap data of ON CELLS
MeansOfOnNormedMeans=[]
PercentileOfOnNormedMeans=[]
OrderedOnReporters=[]
OrderedOnDoses=[]
for reporter in ReporterList:
    data=BSOnNormedMeans[BSOnNormedMeans['Reporter']==reporter]
    for dose in DoseList:
        dosedata=data[data['Dose']==dose]
        for column in ColumnNames:
            iteration=dosedata[column]
            meanofmeans=np.average(iteration)
            percentileofmeans=np.percentile(iteration,90)
            OrderedOnReporters.append(reporter)
            OrderedOnDoses.append(dose)
            MeansOfOnNormedMeans.append(meanofmeans)
            PercentileOfOnNormedMeans.append(percentileofmeans)
BootStrapOfOnNormedMeans=pd.DataFrame()
BootStrapOfOnNormedMeans['Reporter']=OrderedOnReporters
BootStrapOfOnNormedMeans['Dose']=OrderedOnDoses
BootStrapOfOnNormedMeans['Mean']=MeansOfOnNormedMeans
BootStrapOfOnNormedMeans['Percentile']=PercentileOfOnNormedMeans

#Work toward getting a bootstrapped ratio of on/off status between DMSO and the U0126 doses, starting by bootstrapping the serial number of cells, contained by dose and reporter
#Use serial number to extract the mode and the experiment number from the corresponding cell in the original data
UOhList=['0.1','0.342']
ProxPOModes=pd.DataFrame()
ProxPTModes=pd.DataFrame()
ProxPOXPs=pd.DataFrame()
ProxPTXPs=pd.DataFrame()
DistalPOModes=pd.DataFrame()
DistalPTModes=pd.DataFrame()
DistalPOXPs=pd.DataFrame()
DistalPTXPs=pd.DataFrame()
FullPOModes=pd.DataFrame()
FullPTModes=pd.DataFrame()
FullPOXPs=pd.DataFrame()
FullPTXPs=pd.DataFrame()
ProxModeDFs=[ProxPOModes,ProxPTModes]
ProxXPDFs=[ProxPOXPs,ProxPTXPs]
DistalModeDFs=[DistalPOModes,DistalPTModes]
DistalXPDFs=[DistalPOXPs,DistalPTXPs]
FullModeDFs=[FullPOModes,FullPTModes]
FullXPDFs=[FullPOXPs,FullPTXPs]
ModeDFs=[ProxModeDFs,DistalModeDFs,FullModeDFs]
XPDFs=[ProxXPDFs,DistalXPDFs,FullXPDFs]
for reporter,modedfs,xpdfs in zip(ReporterList,ModeDFs,XPDFs):
    data=Measurements[Measurements['Reporter']==reporter]
    for dose,modedf,xpdf in zip(UOhList,modedfs,xpdfs):
        dosedata=data[data['Dose']==dose]
        serial=dosedata.SerialNum.tolist()
        bsserial=np.random.choice(serial,(len(serial),1000))
        for column,name in zip(range(1000),ColumnNames):
            replicate=bsserial[:,column]
            modelist=[]
            xplist=[]
            for cell in range(len(replicate)):
                index=replicate[cell]-1
                mode=Measurements.Mode[index]
                xp=Measurements['Experiment Number'][index]
                modelist.append(mode)
                xplist.append(xp)
            modedf[name]=modelist
            xpdf[name]=xplist
        xpdf.to_csv('FINAL VERSION Bootstrapped XP Table for '+reporter+' '+dose+'.csv')
        modedf.to_csv('FINAL VERSION Bootstrapped Mode Table for '+reporter+' '+dose+'.csv')
            
#Use the DFs containing the experiment numbers to bootstrap equal amounts of DMSO embryos for each bootstrap replicate
#This must be done once for each U0126 treatment group, generating TWO DMSO controls per reporter
ProxDMSOPO=pd.DataFrame()
DistalDMSOPO=pd.DataFrame()
FullDMSOPO=pd.DataFrame()
ProxDMSOPT=pd.DataFrame()
DistalDMSOPT=pd.DataFrame()
FullDMSOPT=pd.DataFrame()
ProxDMSO=[ProxDMSOPO,ProxDMSOPT]
DistalDMSO=[DistalDMSOPO,DistalDMSOPT]
FullDMSO=[FullDMSOPO,FullDMSOPT]
DMSODFs=[ProxDMSO,DistalDMSO,FullDMSO]
for reporter,xpdf,dmso in zip(ReporterList,XPDFs,DMSODFs):
    data=Measurements[Measurements['Reporter']==reporter]
    controldata=data[data['Dose']=='DMSO']
    for dose,dmsodose,dosename in zip(xpdf,dmso,UOhList):
        for column in ColumnNames:
            reptomatch=dose[column].tolist()
            itercontrolboot=[]
            for xp in XPList:
                count=reptomatch.count(xp)
                if count>0:
                    datatoboot=controldata[controldata['Experiment Number']==xp]
                    datatoboot=datatoboot['Mode'].tolist()
                    controlboot=np.random.choice(datatoboot,count)
                    itercontrolboot=np.concatenate((itercontrolboot,controlboot))
            dmsodose[column]=itercontrolboot
        dmsodose.to_csv('FINAL VERSION Bootstrapped Control Modes for '+reporter+' '+dosename+'.csv')

#Generate a list of U0126 proportion on / DMSO proportion on for each reporter/dose combo
#This ratio directly corresponds to the remaining fraction of cells that still have expression, following treatment, relative to DMSO (and controlled for electroporation efficiency)
PropDiffs=pd.DataFrame()
PropDiffs['Bootstrap Replicate']=ColumnNames
for reporter,modedf,dmso in zip(ReporterList,ModeDFs,DMSODFs):
    for dose,dmsomatch,dosename in zip(modedf,dmso,UOhList):
        propdiffs=[]
        for column in ColumnNames:
            uohcolumn=dose[column]
            uohcount=sum(i>10.5 for i in uohcolumn)
            uohpropon=uohcount/len(uohcolumn)
            dmsocolumn=dmsomatch[column]
            dmsocount=sum(i>10.5 for i in dmsocolumn)
            dmsopropon=dmsocount/len(dmsocolumn)
            propdiff=uohpropon/dmsopropon
            propdiffs.append(propdiff)
        PropDiffs[reporter+dosename]=propdiffs

#Run stats (p-val, 95% Bootstrap CI) on each on/off distribution, comparing the distributions to one
PropDiffNames=['Prox0.1','Prox0.342','Distal0.1','Distal0.342','FL0.1','FL0.342']
TList=[]
PList=[]
CILowerList=[]
CIUpperList=[]
MeanList=[]
for name in PropDiffNames:
    data=PropDiffs[name]
    t,p=ttest_1samp(data,1)
    TList.append(t)
    PList.append(p)
    cilower,ciupper=np.percentile(data,[2.5,97.5])
    mean=np.average(data)
    CILowerList.append(cilower)
    CIUpperList.append(ciupper)
    MeanList.append(mean)
TTestResults=pd.DataFrame()
TTestResults['ComparisonGroup']=PropDiffNames
TTestResults['TStatistics']=TList
TTestResults['PValues']=PList
TTestResults['95%CILowerBound']=CILowerList
TTestResults['Mean']=MeanList
TTestResults['95%CIUpperBound']=CIUpperList
TTestResults.to_csv('FINAL VERSION Bootstrapped OnOff Proportion T test Results.csv')

#Make a complete table of bootstrap 95% confidence intervals (with medians) for the expression ratio, mean of normed means, and percentile of normed means (normed means are for ON CELLS)
MeasureList=[]
CIReporterList=[]
CIDoseList=[]
CILowerList=[]
MedianList=[]
CIUpperList=[]
for name in PropDiffNames:
    data=PropDiffs[name]
    cilower,ciupper=np.percentile(data,[2.5,97.5])
    median=np.median(data)
    if name[0]=='P':
        reporter='Proximal'
    elif name[0]=='D':
        reporter='Distal'
    else:
        reporter='Full Length'
    if name[-1]=='1':
        dose=0.1
    else:
        dose=0.342
    MeasureList.append('Ratio')
    CIReporterList.append(reporter)
    CIDoseList.append(dose)
    CILowerList.append(cilower)
    CIUpperList.append(ciupper)
    MedianList.append(median)
OfMeansList=['Mean of Normalized Means','90th Percentile of Normalized Means']
VarList=['Mean','Percentile']
for measure,var in zip(OfMeansList,VarList):
    for reporter in ReporterList:
        data=BootStrapOfOnNormedMeans[BootStrapOfOnNormedMeans['Reporter']==reporter]
        for dose in DoseList:
            dosedata=data[data['Dose']==dose]
            dosedata=dosedata[var]
            cilower,ciupper=np.percentile(dosedata,[2.5,97.5])
            median=np.median(dosedata)
            MeasureList.append(measure)
            CIReporterList.append(reporter)
            CIDoseList.append(dose)
            CILowerList.append(cilower)
            CIUpperList.append(ciupper)
            MedianList.append(median)
BootCIs=pd.DataFrame()
BootCIs['Measurement']=MeasureList
BootCIs['Reporter']=CIReporterList
BootCIs['Dose']=CIDoseList
BootCIs['95% Confidence Interval Lower Bound']=CILowerList
BootCIs['Median']=MedianList
BootCIs['95% Confidence Interval Upper Bound']=CIUpperList
BootCIs.to_csv('FINAL VERSION Bootstrap Confidence Intervals Table.csv')

#Make barplots of the proportion on, mean of means, and mean of 90th percentiles, using bootstrap CIs as the error bars
OnOffBS=BootCIs[BootCIs['Measurement']=='Ratio']
RatioPOs=OnOffBS[OnOffBS['Dose']=='0.1']
RatioPTs=OnOffBS[OnOffBS['Dose']=='0.342']
RatioPOLower=RatioPOs['Median']-RatioPOs['95% Confidence Interval Lower Bound']
RatioPOUpper=RatioPOs['95% Confidence Interval Upper Bound']-RatioPOs['Median']
RatioPTLower=RatioPTs['Median']-RatioPTs['95% Confidence Interval Lower Bound']
RatioPTUpper=RatioPTs['95% Confidence Interval Upper Bound']-RatioPTs['Median']
fig,ax=plt.subplots(figsize=(8,6))
POs=ax.bar([1-0.175,2-0.175,3-0.175],RatioPOs['Median'],0.35,yerr=[RatioPOLower,RatioPOUpper],label='0.1 uM')
PTs=ax.bar([1+0.175,2+0.175,3+0.175],RatioPTs['Median'],0.35,yerr=[RatioPTLower,RatioPTUpper],label='0.342 uM')
fig=plt.hlines(1,0.5,3.5)
plt.setp(ax,xlabel='Reporter',xticks=[1,2,3],xticklabels=['Proximal','Distal','Full Length'],xlim=(0.5,3.5),ylim=(0,1.2),ylabel='Proportion on relative to DMSO')
plt.legend()
fig.figure.savefig('FINAL VERSION On Off Bootstrap Barplot.pdf',transparent=True)

MOMsBS=BootCIs[BootCIs['Measurement']=='Mean of Normalized Means']
MOMsC=MOMsBS[MOMsBS['Dose']=='DMSO']
MOMsPO=MOMsBS[MOMsBS['Dose']=='0.1']
MOMsPT=MOMsBS[MOMsBS['Dose']=='0.342']
MOMsCLower=MOMsC['Median']-MOMsC['95% Confidence Interval Lower Bound']
MOMsCUpper=MOMsC['95% Confidence Interval Upper Bound']-MOMsC['Median']
MOMsPOLower=MOMsPO['Median']-MOMsPO['95% Confidence Interval Lower Bound']
MOMsPOUpper=MOMsPO['95% Confidence Interval Upper Bound']-MOMsPO['Median']
MOMsPTLower=MOMsPT['Median']-MOMsPT['95% Confidence Interval Lower Bound']
MOMsPTUpper=MOMsPT['95% Confidence Interval Upper Bound']-MOMsPT['Median']
fig,ax=plt.subplots(figsize=(8,6))
Cs=ax.bar([1-.25,2-.25,3-.25],MOMsC['Median'],0.25,yerr=[MOMsCLower,MOMsCUpper],label='DMSO')
POs=ax.bar([1,2,3],MOMsPO['Median'],0.25,yerr=[MOMsPOLower,MOMsPOUpper],label='0.1 uM')
PTs=ax.bar([1+.25,2+.25,3+.25],MOMsPT['Median'],0.25,yerr=[MOMsPTLower,MOMsPTUpper],label='0.342 uM')
plt.setp(ax,xlabel='Reporter',xticks=[1,2,3],xticklabels=['Proximal','Distal','Full Length'],xlim=(0.5,3.5),ylim=(0,1.5),ylabel='Mean of Means')
plt.legend()
fig.savefig('FINAL VERSION Mean of Means Barplot.pdf',transparent=True)

POMsBS=BootCIs[BootCIs['Measurement']=='90th Percentile of Normalized Means']
POMsC=POMsBS[POMsBS['Dose']=='DMSO']
POMsPO=POMsBS[POMsBS['Dose']=='0.1']
POMsPT=POMsBS[POMsBS['Dose']=='0.342']
POMsCLower=POMsC['Median']-POMsC['95% Confidence Interval Lower Bound']
POMsCUpper=POMsC['95% Confidence Interval Upper Bound']-POMsC['Median']
POMsPOLower=POMsPO['Median']-POMsPO['95% Confidence Interval Lower Bound']
POMsPOUpper=POMsPO['95% Confidence Interval Upper Bound']-POMsPO['Median']
POMsPTLower=POMsPT['Median']-POMsPT['95% Confidence Interval Lower Bound']
POMsPTUpper=POMsPT['95% Confidence Interval Upper Bound']-POMsPT['Median']
fig,ax=plt.subplots(figsize=(8,6))
Cs=ax.bar([1-.25,2-.25,3-.25],POMsC['Median'],0.25,yerr=[POMsCLower,POMsCUpper],label='DMSO')
POs=ax.bar([1,2,3],POMsPO['Median'],0.25,yerr=[POMsPOLower,POMsPOUpper],label='0.1 uM')
PTs=ax.bar([1+.25,2+.25,3+.25],POMsPT['Median'],0.25,yerr=[POMsPTLower,POMsPTUpper],label='0.342 uM')
plt.setp(ax,xlabel='Reporter',xticks=[1,2,3],xticklabels=['Proximal','Distal','Full Length'],xlim=(0.5,3.5),ylabel='90th Percentile of Means')
plt.legend()
fig.savefig('FINAL VERSION 90th Percentile of Means Barplot.pdf',transparent=True)

#Produce plots of histogram distributions of just the normalized mean, with all three doses plotted as individual lines for each reporter
#Do this for both ALL cells (with no adjustment) and ON cells (adjusting each histogram bin for the proportion on)
ProxOn=OnCells[OnCells['Reporter']=='Prox']
DistalOn=OnCells[OnCells['Reporter']=='Distal']
FullOn=OnCells[OnCells['Reporter']=='FL']
ProxOnControl=ProxOn[ProxOn['Dose']=='DMSO']
DistalOnControl=DistalOn[DistalOn['Dose']=='DMSO']
FullOnControl=FullOn[FullOn['Dose']=='DMSO']
ProxOnPointOne=ProxOn[ProxOn['Dose']=='0.1']
DistalOnPointOne=DistalOn[DistalOn['Dose']=='0.1']
FullOnPointOne=FullOn[FullOn['Dose']=='0.1']
ProxOnPointThree=ProxOn[ProxOn['Dose']=='0.342']
DistalOnPointThree=DistalOn[DistalOn['Dose']=='0.342']
FullOnPointThree=FullOn[FullOn['Dose']=='0.342']
ReporterOnData=[ProxOn,DistalOn,FullOn]

ProxAll=Measurements[Measurements['Reporter']=='Prox']
DistalAll=Measurements[Measurements['Reporter']=='Distal']
FLAll=Measurements[Measurements['Reporter']=='FL']
ProxAllControl=ProxAll[ProxAll['Dose']=='DMSO']
ProxAllPointOne=ProxAll[ProxAll['Dose']=='0.1']
ProxAllPointThree=ProxAll[ProxAll['Dose']=='0.342']
DistalAllControl=DistalAll[DistalAll['Dose']=='DMSO']
DistalAllPointOne=DistalAll[DistalAll['Dose']=='0.1']
DistalAllPointThree=DistalAll[DistalAll['Dose']=='0.342']
FLAllControl=FLAll[FLAll['Dose']=='DMSO']
FLAllPointOne=FLAll[FLAll['Dose']=='0.1']
FLAllPointThree=FLAll[FLAll['Dose']=='0.342']

ConditionOnList=[ProxOnControl,DistalOnControl,FullOnControl,ProxOnPointOne,DistalOnPointOne,FullOnPointOne,ProxOnPointThree,DistalOnPointThree,FullOnPointThree]
ConditionAllList=[ProxAllControl,DistalAllControl,FLAllControl,ProxAllPointOne,DistalAllPointOne,FLAllPointOne,ProxAllPointThree,DistalAllPointThree,FLAllPointThree]

MeanhistlistOn=[]
MeanhistlistAll=[]
controlline=mlines.Line2D([],[],color='b',linestyle='-',label='DMSO')
poline=mlines.Line2D([],[],color='orange',linestyle='-',label='0.1 uM')
ptline=mlines.Line2D([],[],color='g',linestyle='-',label='0.342 uM')
for on,allcells in zip(ConditionOnList,ConditionAllList):
    histogramon=plt.hist(np.log2(on['XPNormedScaledMean']+1),bins=meanbins,density=True)
    histogramall=plt.hist(np.log2(allcells['XPNormedScaledMean']+1),bins=meanbins,density=True)
    MeanhistlistOn.append(histogramon[0])
    MeanhistlistAll.append(histogramall[0])
plt.clf()

TrueMeanhistlistOn=[]
for onhist,counter in zip(MeanhistlistOn,range(len(MeanhistlistOn))):
    if counter==3:
        onhist=onhist*OnOffBS['Median'][0]
    if counter==4:
        onhist=onhist*OnOffBS['Median'][2]
    if counter==5:
        onhist=onhist*OnOffBS['Median'][4]
    if counter==6:
        onhist=onhist*OnOffBS['Median'][1]
    if counter==7:
        onhist=onhist*OnOffBS['Median'][3]
    if counter==8:
        onhist=onhist*OnOffBS['Median'][5]
    TrueMeanhistlistOn.append(onhist)

fig,ax=plt.subplots(figsize=(8,6))
ax.plot(meanx,TrueMeanhistlistOn[0],'b')
ax.plot(meanx,TrueMeanhistlistOn[3],'orange')
ax.plot(meanx,TrueMeanhistlistOn[6],'g')
plt.setp(ax,title='Proximal',ylabel='Adjusted Bin Density',ylim=(-0.01,0.5),xlabel='Mean Expression')
plt.legend(handles=[controlline,poline,ptline],loc='upper right')
fig.savefig('FINAL VERSION ON CELLS Normalized Means Proximal Distributions.pdf',transparent=True)

fig,ax=plt.subplots(figsize=(8,6))
ax.plot(meanx,TrueMeanhistlistOn[1],'b')
ax.plot(meanx,TrueMeanhistlistOn[4],'orange')
ax.plot(meanx,TrueMeanhistlistOn[7],'g')
plt.setp(ax,title='Distal',ylabel='Adjusted Bin Density',ylim=(-0.01,0.5),xlabel='Mean Expression')
plt.legend(handles=[controlline,poline,ptline],loc='upper right')
fig.savefig('FINAL VERSION ON CELLS Normalized Means Distal Distributions.pdf',transparent=True)

fig,ax=plt.subplots(figsize=(8,6))
ax.plot(meanx,TrueMeanhistlistOn[2],'b')
ax.plot(meanx,TrueMeanhistlistOn[5],'orange')
ax.plot(meanx,TrueMeanhistlistOn[8],'g')
plt.setp(ax,title='Full Length',ylabel='Adjusted Bin Density',ylim=(-0.01,0.5),xlabel='Mean Expression')
plt.legend(handles=[controlline,poline,ptline],loc='upper right')
fig.savefig('FINAL VERSION ON CELLS Normalized Means Full Length Distributions.pdf',transparent=True)

fig,ax=plt.subplots(figsize=(8,6))
ax.plot(meanx,MeanhistlistAll[0],'b')
ax.plot(meanx,MeanhistlistAll[3],'orange')
ax.plot(meanx,MeanhistlistAll[6],'g')
plt.setp(ax,title='Proximal',ylabel='Bin Density',ylim=(-0.01,0.5),xlabel='Mean Expression')
plt.legend(handles=[controlline,poline,ptline],loc='upper right')
fig.savefig('FINAL VERSION ALL CELLS Normalized Means Proximal Distributions.pdf',transparent=True)

fig,ax=plt.subplots(figsize=(8,6))
ax.plot(meanx,MeanhistlistAll[1],'b')
ax.plot(meanx,MeanhistlistAll[4],'orange')
ax.plot(meanx,MeanhistlistAll[7],'g')
plt.setp(ax,title='Distal',ylabel='Bin Density',ylim=(-0.01,0.5),xlabel='Mean Expression')
plt.legend(handles=[controlline,poline,ptline],loc='upper right')
fig.savefig('FINAL VERSION ALL CELLS Normalized Means Distal Distributions.pdf',transparent=True)

fig,ax=plt.subplots(figsize=(8,6))
ax.plot(meanx,MeanhistlistAll[2],'b')
ax.plot(meanx,MeanhistlistAll[5],'orange')
ax.plot(meanx,MeanhistlistAll[8],'g')
plt.setp(ax,title='Full Length',ylabel='Bin Density',ylim=(-0.01,0.5),xlabel='Mean Expression')
plt.legend(handles=[controlline,poline,ptline],loc='upper right')
fig.savefig('FINAL VERSION ALL CELLS Normalized Means Full Length Distributions.pdf',transparent=True)

#Take differences between bootstrap populations to get p-values for comparisons in the bar plots

PControl=pd.DataFrame()
PPO=pd.DataFrame()
PPT=pd.DataFrame()
DControl=pd.DataFrame()
DPO=pd.DataFrame()
DPT=pd.DataFrame()
FControl=pd.DataFrame()
FPO=pd.DataFrame()
FPT=pd.DataFrame()

PDFs=[PControl,PPO,PPT]
DDFs=[DControl,DPO,DPT]
FDFs=[FControl,FPO,FPT]
ReporterDFs=[PDFs,DDFs,FDFs]

for reporter,destination in zip(ReporterList,ReporterDFs):
    allmeans=BootStrapOfNormedMeans[BootStrapOfNormedMeans['Reporter']==reporter]
    onmeans=BootStrapOfOnNormedMeans[BootStrapOfOnNormedMeans['Reporter']==reporter]
    for dose,dosedf in zip(DoseList,destination):
        doseall=allmeans[allmeans['Dose']==dose]
        doseon=onmeans[onmeans['Dose']==dose]
        doseall=doseall.Mean.tolist()
        doseonmean=doseon.Mean.tolist()
        doseonperc=doseon.Percentile.tolist()
        dosedf['AllMean']=doseall
        dosedf['OnMean']=doseonmean
        dosedf['OnPerc']=doseonperc

CompareList=[[0,1],[0,2],[1,2]]
PPVals=pd.DataFrame()
DPVals=pd.DataFrame()
FPVals=pd.DataFrame()
ReporterPVals=[PPVals,DPVals,FPVals]

for reporter,df,pvals in zip(ReporterList,ReporterDFs,ReporterPVals):
    allmeanps=[]
    onmeanps=[]
    onpercps=[]
    for compare in CompareList:
        allmeanp=0
        onmeanp=0
        onpercp=0
        groupone=df[compare[0]]
        grouptwo=df[compare[1]]
        grouponeallmean=groupone['AllMean'].tolist()
        grouponeonmean=groupone['OnMean'].tolist()
        grouponeonperc=groupone['OnPerc'].tolist()
        grouptwoallmean=grouptwo['AllMean'].tolist()
        grouptwoonmean=grouptwo['OnMean'].tolist()
        grouptwoonperc=grouptwo['OnPerc'].tolist()
        for one,two in zip(grouponeallmean,grouptwoallmean):
            diff=one-two
            if diff<=0:
                allmeanp=allmeanp+0.001
        for one,two in zip(grouponeonmean,grouptwoonmean):
            diff=one-two
            if diff<=0:
                onmeanp=onmeanp+0.001
        for one,two in zip(grouponeonperc,grouptwoonperc):
            diff=one-two
            if diff<=0:
                onpercp=onpercp+0.001
        allmeanps.append(allmeanp)
        onmeanps.append(onmeanp)
        onpercps.append(onpercp)
    pvals['Comparison']=['DMSO to 0.1','DMSO to 0.342','0.1 to 0.342']
    pvals['AllMeans']=allmeanps
    pvals['OnMeans']=onmeanps
    pvals['OnPercs']=onpercps
        
#Make a barplot of the mean ALL cell expression, using bootstrap CIs as the error bars
ALLC=[np.median(PControl.AllMean.tolist()),np.median(DControl.AllMean.tolist()),np.median(FControl.AllMean.tolist())]
ALLPO=[np.median(PPO.AllMean.tolist()),np.median(DPO.AllMean.tolist()),np.median(FPO.AllMean.tolist())]
ALLPT=[np.median(PPT.AllMean.tolist()),np.median(DPT.AllMean.tolist()),np.median(FPT.AllMean.tolist())]
ALLCLower=[np.percentile(PControl.AllMean.tolist(),2.5),np.percentile(DControl.AllMean.tolist(),2.5),np.percentile(FControl.AllMean.tolist(),2.5)]
ALLCLower=[ALLC[0]-ALLCLower[0],ALLC[1]-ALLCLower[1],ALLC[2]-ALLCLower[2]]
ALLCUpper=[np.percentile(PControl.AllMean.tolist(),97.5),np.percentile(DControl.AllMean.tolist(),97.5),np.percentile(FControl.AllMean.tolist(),97.5)]
ALLCUpper=[ALLCUpper[0]-ALLC[0],ALLCUpper[1]-ALLC[1],ALLCUpper[2]-ALLC[2]]
ALLPOLower=[np.percentile(PPO.AllMean.tolist(),2.5),np.percentile(DPO.AllMean.tolist(),2.5),np.percentile(FPO.AllMean.tolist(),2.5)]
ALLPOLower=[ALLPO[0]-ALLPOLower[0],ALLPO[1]-ALLPOLower[1],ALLPO[2]-ALLPOLower[2]]
ALLPOUpper=[np.percentile(PPO.AllMean.tolist(),97.5),np.percentile(DPO.AllMean.tolist(),97.5),np.percentile(FPO.AllMean.tolist(),97.5)]
ALLPOUpper=[ALLPOUpper[0]-ALLPO[0],ALLPOUpper[1]-ALLPO[1],ALLPOUpper[2]-ALLPO[2]]
ALLPTLower=[np.percentile(PPT.AllMean.tolist(),2.5),np.percentile(DPT.AllMean.tolist(),2.5),np.percentile(FPT.AllMean.tolist(),2.5)]
ALLPTLower=[ALLPT[0]-ALLPTLower[0],ALLPT[1]-ALLPTLower[1],ALLPT[2]-ALLPTLower[2]]
ALLPTUpper=[np.percentile(PPT.AllMean.tolist(),97.5),np.percentile(DPT.AllMean.tolist(),97.5),np.percentile(FPT.AllMean.tolist(),97.5)]
ALLPTUpper=[ALLPTUpper[0]-ALLPT[0],ALLPTUpper[1]-ALLPT[1],ALLPTUpper[2]-ALLPT[2]]
fig,ax=plt.subplots(figsize=(8,6))
Cs=ax.bar([1-.25,2-.25,3-.25],ALLC,0.25,yerr=[ALLCLower,ALLCUpper],label='DMSO')
POs=ax.bar([1,2,3],ALLPO,0.25,yerr=[ALLPOLower,ALLPOUpper],label='0.1 uM')
PTs=ax.bar([1+.25,2+.25,3+.25],ALLPT,0.25,yerr=[ALLPTLower,ALLPTUpper],label='0.342 uM')
plt.setp(ax,xlabel='Reporter',xticks=[1,2,3],xticklabels=['Proximal','Distal','Full Length'],xlim=(0.5,3.5),ylim=(0,1.2),ylabel='Mean of Means')
plt.legend()
fig.savefig('FINAL VERSION Mean of ALL Means Barplot.pdf',transparent=True)
```
