## Supplemental Appendix S3 for "*Ciona Brachyury* proximal and distal enhancers have different FGF dose-response relationships": Fig 4 and Associated Supplement.py

```
#!/usr/bin/env python3
# -*- coding: utf-8 -*-
"""
Created on Fri May 22 11:09:02 2020

@author: Matt
"""

#Import necessary libraries
import numpy as np
import pandas as pd
import matplotlib.pyplot as plt
import seaborn as sns
import matplotlib.colors as mcolors
import random
import os
import matplotlib.lines as mlines
from scipy.stats import f_oneway
from statsmodels.stats.multicomp import MultiComparison
plt.rcParams['pdf.fonttype']=42

#Change into input/output directory, read in data, and make dose adjustments for DMSO
os.chdir("/Volumes/MH Veeman Lab Home Backup/in situ")
InSitu=pd.read_csv('InSituForPython.csv')
SEVData=pd.read_csv('SEVData.csv')
InSituNonUniform=pd.read_csv('InSituNonUniform.csv')
FLData=SEVData[SEVData['enhancer_construct']=='Shadow+Spacer+Proximal']
AveragedScores=InSitu.iloc[:,25:]
Doses=InSitu['LogDose']
LogDoses=np.log10(Doses)
AveragedScores['LogDose']=LogDoses
AdjustedDoses=[]
for dose in LogDoses:
    if dose<=(-1.3):
        altereddose=np.log10(1/64)
    else:
        altereddose=dose
    AdjustedDoses.append(altereddose)
AveragedScores['ModLogDose']=AdjustedDoses

#Melt data to be easier to work with, split into subdivisions by cell pair
MeltedAverage=pd.melt(AveragedScores,id_vars=["DoseCode","LogDose",'ModLogDose'],value_vars=["Secondary","A8.14","A8.13","A8.6","A8.5"],var_name="Cell Pair",value_name="Score")
Secondary=pd.melt(InSitu,id_vars=['LogDose'],value_vars=['Secondary'],var_name='Cell Pair',value_name="Score")
Secondary['LogDose']=np.log10(Secondary['LogDose'])
SecondaryDoses=Secondary['LogDose']
SecondaryScores=Secondary['Score']
SecN=len(Secondary)
DCells=pd.melt(InSitu,id_vars=['LogDose'],value_vars=['A8.14'],var_name='Cell Pair',value_name="Score")
DCells['LogDose']=np.log10(DCells['LogDose'])
DDoses=DCells['LogDose']
DScores=DCells['Score']
DN=len(DCells)
CCells=pd.melt(InSitu,id_vars=['LogDose'],value_vars=['A8.13'],var_name='Cell Pair',value_name="Score")
CCells['LogDose']=np.log10(CCells['LogDose'])
CDoses=CCells['LogDose']
CScores=CCells['Score']
CN=len(CCells)
BCells=pd.melt(InSitu,id_vars=['LogDose'],value_vars=['A8.6'],var_name='Cell Pair',value_name="Score")
BCells['LogDose']=np.log10(BCells['LogDose'])
BDoses=BCells['LogDose']
BScores=BCells['Score']
BN=len(BCells)
ACells=pd.melt(InSitu,id_vars=['LogDose'],value_vars=['A8.5'],var_name='Cell Pair',value_name="Score")
ACells['LogDose']=np.log10(ACells['LogDose'])
ADoses=ACells['LogDose']
AScores=ACells['Score']
AN=len(ACells)

#Use cell pair scores to generate average scores over whole embryos
InSituAverage=[]
for a,b,c,d,sec in zip(AScores,BScores,CScores,DScores,SecondaryScores):
    total=np.average((a,b,c,d,sec))
    InSituAverage.append(total)
InSituSingle=pd.DataFrame()
InSituSingle['OriginalDoses']=ADoses
InSituSingle['AverageExpression']=InSituAverage
SEVAdjustedDoses=[]
for dose in ADoses:
    if dose<=(-1.3):
        altereddose=np.log10(0.00855)
    else:
        altereddose=dose
    SEVAdjustedDoses.append(altereddose)
InSituSingle['AdjustedDose']=SEVAdjustedDoses

#Rescale whole-embryo averages to have a common DMSO score with the reporter data
DMSODose=InSituSingle[InSituSingle['AdjustedDose']<=(-1.3)]
ScalingFactor=1/(np.average(DMSODose.AverageExpression))
ScaledExpression=[]
for score in InSituAverage:
    scaledscore=score*ScalingFactor
    ScaledExpression.append(scaledscore)
InSituSingle['ScaledExpression']=ScaledExpression

#Add a column to combined in situ data for proper plotting
AllCellsColumn=[]
for embryo in range(len(InSituSingle)):
    AllCellsColumn.append('combined')
InSituSingle['AllCellsColumn']=AllCellsColumn

#Enter basic figure information used by mulitple panels
norm=mcolors.Normalize(vmin=0.,vmax=3.)
ticklabels=[' L2',' LD',' LC',' LB',' LA',' RA',' RB',' RC',' RD',' R2']
AVGticklabels=['       2','       D','       C','       B','       A']
pc_kwargs={'rasterized':True,'cmap':'Blues','norm':norm}
detailedx=np.linspace(-1.51,0.61,100)
doseticks=[np.log10(0.015625),np.log10(0.0625),np.log10(0.125),np.log10(0.25),np.log10(0.5),np.log10(1),np.log10(2),np.log10(4)]
doseticklabels=['DMSO','1/16','1/8','1/4','1/2','1','2','4']

#Make plot outlining average scores for each cell type
CellScores,ax=plt.subplots(figsize=(8,6))
CellScores=sns.lineplot(x="ModLogDose",y="Score",hue="Cell Pair",hue_order=['A8.5','A8.6','A8.13','A8.14','Secondary'],ax=ax,ci=95,markers=['o','o','o','o','o'],dashes=False,style='Cell Pair',data=MeltedAverage)
plt.setp(ax,xlabel='[U0126 (uM)]',xticks=doseticks,xticklabels=doseticklabels,ylabel='Mean Expression Score',title='Individual Cell Pairs')
CellScores.figure.savefig('FINAL VERSION in situ All Cell Pair Scores.pdf',transparent=True)

#Make a plot that puts grand average in situ data on shared axes with the FL reporter data
isline=mlines.Line2D([],[],color='purple',label='Average Scaled in situ Expression')
fline=mlines.Line2D([],[],color='g',label='Full Length SEV Expression')
fig,ax=plt.subplots(figsize=(8,6))
fig=sns.lineplot(x='ModLogDose',y='NSSEVNormedToFL0.01581',color='g',style='enhancer_construct',ax=ax,ci=95,markers='o',dashes=False,data=FLData)
fig=sns.lineplot(x='AdjustedDose',y='ScaledExpression',color='purple',style='AllCellsColumn',ax=ax,ci=95,markers='o',dashes=False,data=InSituSingle)
plt.setp(ax,xlabel='[U0126 (uM)]',xticks=[np.log10(0.00855),np.log10(0.0625),np.log10(0.125),np.log10(0.25),np.log10(0.5),np.log10(1),np.log10(2),np.log10(4)],xticklabels=doseticklabels,ylabel='Mean Expression Score',title='in situ and reporter comparison')
plt.legend(handles=[fline,isline],loc='upper right')
fig.figure.savefig('FINAL VERSION SEV and in situ scores comparison.pdf',transparent=True)

#Collapse the medial pairs into one line, lateral pairs into another, and plot both with secondary
MedialAverage=[]
LateralAverage=[]
for a,b,c,d in zip(AScores,BScores,CScores,DScores):
    medial=np.average((a,b))
    lateral=np.average((c,d))
    MedialAverage.append(medial)
    LateralAverage.append(lateral)
ISCombined=pd.DataFrame()
ISCombined['ModLogDose']=AdjustedDoses
ISCombined['MedialCells']=MedialAverage
ISCombined['LateralCells']=LateralAverage
ISCombined['SecondaryCells']=SecondaryScores
MeltedCombined=pd.melt(ISCombined,id_vars=['ModLogDose'],value_vars=["SecondaryCells","LateralCells","MedialCells"],var_name="Cell Type",value_name="Score")
medline=mlines.Line2D([],[],color='b',label='Medial Cells')
latline=mlines.Line2D([],[],color='orange',label='Lateral Cells')
secline=mlines.Line2D([],[],color='g',label='Secondary Cells')
fig,ax=plt.subplots(figsize=(8,6))
fig=sns.lineplot(x="ModLogDose",y="Score",hue="Cell Type",style='Cell Type',hue_order=['MedialCells','LateralCells','SecondaryCells'],palette=['b','orange','g'],ax=ax,ci=95,markers=['o','o','o'],dashes=False,data=MeltedCombined)
plt.setp(ax,xlabel='Log[U0126 (uM)]',xticks=doseticks,xticklabels=doseticklabels,ylabel='Mean Expression Score',title='Combined Cell Types')
plt.legend(handles=[medline,latline,secline],loc='upper right')
fig.figure.savefig('FINAL VERSION in situ Combined Type Cell Scores.pdf',transparent=True)

#Create a clustered bar plot looking at the 3 critical doses (1/4, 1/2, and 1 uM) to see how the cell pairs are different
BarScores=MeltedAverage[MeltedAverage['ModLogDose']<0.1]
BarScores=BarScores[BarScores['ModLogDose']>-0.75]
fig,ax=plt.subplots(figsize=(8,6))
fig=sns.barplot(x='ModLogDose',y='Score',hue='Cell Pair',hue_order=['A8.5','A8.6','A8.13','A8.14','Secondary'],ax=ax,ci=95,data=BarScores)
ax.set_xticklabels(['0.25','0.5','1.0'])
plt.setp(ax,ylabel='Mean Expression Score',xlabel='[U0126 (uM)]')
fig.figure.savefig('FINAL VERSION in situ Critical Doses Barplot.pdf',transparent=True)

#Create a bar plot for just the 1/2 uM dose
ShortBar=MeltedAverage[MeltedAverage['ModLogDose']<0]
ShortBar=ShortBar[ShortBar['ModLogDose']>-0.5]
fig,ax=plt.subplots(figsize=(8,6))
fig=sns.barplot(x='Cell Pair',y='Score',order=['A8.5','A8.6','A8.13','A8.14','Secondary'],ax=ax,ci=95,data=ShortBar)
plt.setp(ax,ylabel='Mean Expression Score',xlabel='Cell Pair')
fig.figure.savefig('FINAL VERSION in situ Half uM Barplot.pdf',transparent=True)

#Run ANOVA and Tukey HSD on the scores at the critical doses to identify cell pairs with differences in behavior
CriticalList=[3,4,5]
DoseNames=['0.25','0.5','1.0']
for dose,dosename in zip(CriticalList,DoseNames):
    flist=[]
    plist=[]
    anovaresults=pd.DataFrame()
    tukeyoutput=pd.DataFrame()
    data=BarScores[BarScores['DoseCode']==dose]
    dataone=data[data['Cell Pair']=='A8.5']
    datatwo=data[data['Cell Pair']=='A8.6']
    datathree=data[data['Cell Pair']=='A8.13']
    datafour=data[data['Cell Pair']=='A8.14']
    datafive=data[data['Cell Pair']=='Secondary']
    f,p=f_oneway(dataone['Score'],datatwo['Score'],datathree['Score'],datafour['Score'],datafive['Score'])
    flist.append(f)
    plist.append(p)
    if p<0.05:
        groupone=dataone['Score']
        grouponelabel=[]
        for cell in groupone:
            grouponelabel.append('A8.5')
        groupone=groupone.to_list()
        grouptwo=datatwo['Score']
        grouptwolabel=[]
        for cell in grouptwo:
            grouptwolabel.append('A8.6')
        grouptwo=grouptwo.to_list()
        groupthree=datathree['Score']
        groupthreelabel=[]
        for cell in groupthree:
            groupthreelabel.append('A8.13')
        groupthree=groupthree.to_list()
        groupfour=datafour['Score']
        groupfourlabel=[]
        for cell in groupfour:
            groupfourlabel.append('A8.14')
        groupfour=groupfour.to_list()
        groupfive=datafive['Score']
        groupfivelabel=[]
        for cell in groupfive:
            groupfivelabel.append('Secondary')
        groupfive=groupfive.to_list()
        groups=groupone+grouptwo+groupthree+groupfour+groupfive
        grouplabels=grouponelabel+grouptwolabel+groupthreelabel+groupfourlabel+groupfivelabel
        mc=MultiComparison(groups,grouplabels)
        result=mc.tukeyhsd()
        tukeyresults=pd.DataFrame(data=result._results_table.data[1:],columns=result._results_table.data[0])
        tukeyoutput=tukeyoutput.append(tukeyresults)
    anovaresults['F-statistic']=flist
    anovaresults['p-value']=plist
    anovaresults.to_csv('FINAL VERSION in situ ANOVA results for '+dosename+' uM.csv')
    tukeyoutput.to_csv('FINAL VERSION in situ Tukey results for '+dosename+' uM.csv')

#Combine ANOVA results into one consolidated table, and Tukey results into another
P25ANOVA=pd.read_csv('FINAL VERSION in situ ANOVA results for 0.25 uM.csv')
P25ANOVA=P25ANOVA.iloc[:,1:]
P5ANOVA=pd.read_csv('FINAL VERSION in situ ANOVA results for 0.5 uM.csv')
P5ANOVA=P5ANOVA.iloc[:,1:]
OANOVA=pd.read_csv('FINAL VERSION in situ ANOVA results for 1.0 uM.csv')
OANOVA=OANOVA.iloc[:,1:]
P25Tukey=pd.read_csv('FINAL VERSION in situ Tukey results for 0.25 uM.csv')
P25Tukey=P25Tukey.iloc[:,1:]
P5Tukey=pd.read_csv('FINAL VERSION in situ Tukey results for 0.5 uM.csv')
P5Tukey=P5Tukey.iloc[:,1:]
OTukey=pd.read_csv('FINAL VERSION in situ Tukey results for 1.0 uM.csv')
OTukey=OTukey.iloc[:,1:]

ANOVACombined=pd.DataFrame(columns=['Dose','F-statistic','p-value'])
ANOVACombined=ANOVACombined.append(P25ANOVA,sort=True)
ANOVACombined=ANOVACombined.append(P5ANOVA,sort=True)
ANOVACombined=ANOVACombined.append(OANOVA,sort=True)
ANOVACombined['Dose']=['0.25 uM','0.5 uM','1.0 uM']

TukeyCombined=pd.DataFrame(columns=['Dose','group1','group2','meandiff','p-adj','lower','upper','reject'])
TukeyCombined=TukeyCombined.append(P25Tukey,sort=True)
TukeyCombined=TukeyCombined.append(P5Tukey,sort=True)
TukeyCombined=TukeyCombined.append(OTukey,sort=True)
TukeyCombined['Dose']=['0.25 uM','0.25 uM','0.25 uM','0.25 uM','0.25 uM','0.25 uM','0.25 uM','0.25 uM','0.25 uM','0.25 uM','0.5 uM','0.5 uM','0.5 uM','0.5 uM','0.5 uM','0.5 uM','0.5 uM','0.5 uM','0.5 uM','0.5 uM','1.0 uM','1.0 uM','1.0 uM','1.0 uM','1.0 uM','1.0 uM','1.0 uM','1.0 uM','1.0 uM','1.0 uM']

ANOVACombined.to_csv('FINAL VERSION in situ full ANOVA results.csv')
TukeyCombined.to_csv('FINAL VERSION in situ full Tukey results.csv')

#Organize data for heatmaps
Scores=np.asarray(InSituNonUniform)
Scores=Scores[:,5:15]
DMSOinsitu=InSituNonUniform[InSituNonUniform['Dose']==0]
DMSOinsitu=np.asarray(DMSOinsitu)
DMSOinsitu=DMSOinsitu[:,5:15]
insitu0p0625=InSituNonUniform[InSituNonUniform['Dose']==0.0625]
insitu0p0625=np.asarray(insitu0p0625)
insitu0p0625=insitu0p0625[:,5:15]
insitu0p125=InSituNonUniform[InSituNonUniform['Dose']==0.125]
insitu0p125=np.asarray(insitu0p125)
insitu0p125=insitu0p125[:,5:15]
insitu0p25=InSituNonUniform[InSituNonUniform['Dose']==0.25]
insitu0p25=np.asarray(insitu0p25)
insitu0p25=insitu0p25[:,5:15]
insitu0p5=InSituNonUniform[InSituNonUniform['Dose']==0.5]
insitu0p5=np.asarray(insitu0p5)
insitu0p5=insitu0p5[:,5:15]
insitu1=InSituNonUniform[InSituNonUniform['Dose']==1]
insitu1=np.asarray(insitu1)
insitu1=insitu1[:,5:15]

#Make heatmaps of cell scores for NONUNIFORM embryos
fullfig,fullax=plt.subplots(figsize=(4,8),constrained_layout=True)
fullim=fullax.pcolormesh(Scores,**pc_kwargs)
fullfig.colorbar(fullim,ax=fullax,shrink=0.6)
plt.tick_params(left="off",labelleft="off")
fullax.set_xticks(np.arange(len(ticklabels)))
fullax.set_xticklabels(ticklabels)
plt.setp(fullax.get_xticklabels(),ha="left")
fullfig.savefig("FINAL VERSION All Doses in situ heatmap.pdf",transparent=True)

DMSOfig,DMSOax=plt.subplots(figsize=(4,8),constrained_layout=True)
DMSOim=DMSOax.pcolormesh(DMSOinsitu,**pc_kwargs)
DMSOfig.colorbar(DMSOim,ax=DMSOax,shrink=0.6)
plt.tick_params(left="off",labelleft="off")
DMSOax.set_xticks(np.arange(len(ticklabels)))
DMSOax.set_xticklabels(ticklabels)
plt.setp(DMSOax.get_xticklabels(),ha="left")
DMSOfig.savefig("FINAL VERSION DMSO in situ heatmap.pdf",transparent=True)

Sixteenthfig,Sixteenthax=plt.subplots(figsize=(4,8),constrained_layout=True)
Sixteenthim=Sixteenthax.pcolormesh(insitu0p0625,**pc_kwargs)
Sixteenthfig.colorbar(Sixteenthim,ax=Sixteenthax,shrink=0.6)
plt.tick_params(left="off",labelleft="off")
Sixteenthax.set_xticks(np.arange(len(ticklabels)))
Sixteenthax.set_xticklabels(ticklabels)
plt.setp(Sixteenthax.get_xticklabels(),ha="left")
Sixteenthfig.savefig("FINAL VERSION 0.0625 uM in situ heatmap.pdf",transparent=True)

Eighthfig,Eighthax=plt.subplots(figsize=(4,8),constrained_layout=True)
Eighthim=Eighthax.pcolormesh(insitu0p125,**pc_kwargs)
Eighthfig.colorbar(Eighthim,ax=Eighthax,shrink=0.6)
plt.tick_params(left="off",labelleft="off")
Eighthax.set_xticks(np.arange(len(ticklabels)))
Eighthax.set_xticklabels(ticklabels)
plt.setp(Eighthax.get_xticklabels(),ha="left")
Eighthfig.savefig("FINAL VERSION 0.125 uM in situ heatmap.pdf",transparent=True)

Fourthfig,Fourthax=plt.subplots(figsize=(4,8),constrained_layout=True)
Fourthim=Fourthax.pcolormesh(insitu0p25,**pc_kwargs)
Fourthfig.colorbar(Fourthim,ax=Fourthax,shrink=0.6)
plt.tick_params(left="off",labelleft="off")
Fourthax.set_xticks(np.arange(len(ticklabels)))
Fourthax.set_xticklabels(ticklabels)
plt.setp(Fourthax.get_xticklabels(),ha="left")
Fourthfig.savefig("FINAL VERSION 0.25 uM in situ heatmap.pdf",transparent=True)

Halffig,Halfax=plt.subplots(figsize=(4,8),constrained_layout=True)
Halfim=Halfax.pcolormesh(insitu0p5,**pc_kwargs)
Halffig.colorbar(Halfim,ax=Halfax,shrink=0.6)
plt.tick_params(left="off",labelleft="off")
Halfax.set_xticks(np.arange(len(ticklabels)))
Halfax.set_xticklabels(ticklabels)
plt.setp(Halfax.get_xticklabels(),ha="left")
Halffig.savefig("FINAL VERSION 0.5 uM in situ heatmap.pdf",transparent=True)

Onefig,Oneax=plt.subplots(figsize=(4,8),constrained_layout=True)
Oneim=Oneax.pcolormesh(insitu1,**pc_kwargs)
Onefig.colorbar(Oneim,ax=Oneax,shrink=0.6)
plt.tick_params(left="off",labelleft="off")
Oneax.set_xticks(np.arange(len(ticklabels)))
Oneax.set_xticklabels(ticklabels)
plt.setp(Oneax.get_xticklabels(),ha="left")
Onefig.savefig("FINAL VERSION 1 uM in situ heatmap.pdf",transparent=True)

#Shuffle cell scores from each embryo into random order
Shuffled=np.zeros((1,10))
RowNumber,ColNumber=np.shape(Scores)
for row in range(0,RowNumber):
    toshuffle=Scores[row,:]
    toshuffle=toshuffle.tolist()
    toshuffle=random.sample(toshuffle,len(toshuffle))
    toshuffle=np.array(toshuffle)
    toshuffle=np.reshape(toshuffle,(1,10))
    Shuffled=np.append(Shuffled,toshuffle,axis=0)
Shuffled=Shuffled[1:,:]

ShuffledDMSO=np.zeros((1,10))
RowNumber,ColNumber=np.shape(DMSOinsitu)
for row in range(0,RowNumber):
    toshuffle=DMSOinsitu[row,:]
    toshuffle=toshuffle.tolist()
    toshuffle=random.sample(toshuffle,len(toshuffle))
    toshuffle=np.array(toshuffle)
    toshuffle=np.reshape(toshuffle,(1,10))
    ShuffledDMSO=np.append(ShuffledDMSO,toshuffle,axis=0)
ShuffledDMSO=ShuffledDMSO[1:,:]

Shuffledsixteenth=np.zeros((1,10))
RowNumber,ColNumber=np.shape(insitu0p0625)
for row in range(0,RowNumber):
    toshuffle=insitu0p0625[row,:]
    toshuffle=toshuffle.tolist()
    toshuffle=random.sample(toshuffle,len(toshuffle))
    toshuffle=np.array(toshuffle)
    toshuffle=np.reshape(toshuffle,(1,10))
    Shuffledsixteenth=np.append(Shuffledsixteenth,toshuffle,axis=0)
Shuffledsixteenth=Shuffledsixteenth[1:,:]

Shuffledeighth=np.zeros((1,10))
RowNumber,ColNumber=np.shape(insitu0p125)
for row in range(0,RowNumber):
    toshuffle=insitu0p125[row,:]
    toshuffle=toshuffle.tolist()
    toshuffle=random.sample(toshuffle,len(toshuffle))
    toshuffle=np.array(toshuffle)
    toshuffle=np.reshape(toshuffle,(1,10))
    Shuffledeighth=np.append(Shuffledeighth,toshuffle,axis=0)
Shuffledeighth=Shuffledeighth[1:,:]

Shuffledfourth=np.zeros((1,10))
RowNumber,ColNumber=np.shape(insitu0p25)
for row in range(0,RowNumber):
    toshuffle=insitu0p25[row,:]
    toshuffle=toshuffle.tolist()
    toshuffle=random.sample(toshuffle,len(toshuffle))
    toshuffle=np.array(toshuffle)
    toshuffle=np.reshape(toshuffle,(1,10))
    Shuffledfourth=np.append(Shuffledfourth,toshuffle,axis=0)
Shuffledfourth=Shuffledfourth[1:,:]

Shuffledhalf=np.zeros((1,10))
RowNumber,ColNumber=np.shape(insitu0p5)
for row in range(0,RowNumber):
    toshuffle=insitu0p5[row,:]
    toshuffle=toshuffle.tolist()
    toshuffle=random.sample(toshuffle,len(toshuffle))
    toshuffle=np.array(toshuffle)
    toshuffle=np.reshape(toshuffle,(1,10))
    Shuffledhalf=np.append(Shuffledhalf,toshuffle,axis=0)
Shuffledhalf=Shuffledhalf[1:,:]

Shuffledone=np.zeros((1,10))
RowNumber,ColNumber=np.shape(insitu1)
for row in range(0,RowNumber):
    toshuffle=insitu1[row,:]
    toshuffle=toshuffle.tolist()
    toshuffle=random.sample(toshuffle,len(toshuffle))
    toshuffle=np.array(toshuffle)
    toshuffle=np.reshape(toshuffle,(1,10))
    Shuffledone=np.append(Shuffledone,toshuffle,axis=0)
Shuffledone=Shuffledone[1:,:]

#Create heatmaps of the random shuffled embryos
randomfig,randomax=plt.subplots(figsize=(4,8),constrained_layout=True)
randomim=randomax.pcolormesh(Shuffled,**pc_kwargs)
randomfig.colorbar(randomim,ax=randomax,shrink=0.6)
plt.tick_params(left="off",labelleft="off")
randomax.set_xticks(np.arange(len(ticklabels)))
randomax.set_xticklabels(ticklabels)
plt.setp(randomax.get_xticklabels(),ha="left")
randomfig.savefig("FINAL VERSION Random order all nonuniform cells in situ heatmap.pdf",transparent=True)

randomDMSOfig,randomDMSOax=plt.subplots(figsize=(4,8),constrained_layout=True)
randomDMSOim=randomDMSOax.pcolormesh(ShuffledDMSO,**pc_kwargs)
randomDMSOfig.colorbar(randomDMSOim,ax=randomDMSOax,shrink=0.6)
plt.tick_params(left="off",labelleft="off")
randomDMSOax.set_xticks(np.arange(len(ticklabels)))
randomDMSOax.set_xticklabels(ticklabels)
plt.setp(randomDMSOax.get_xticklabels(),ha="left")
randomDMSOfig.savefig("FINAL VERSION Random order DMSO in situ heatmap.pdf",transparent=True)

randomsixteenthfig,randomsixteenthax=plt.subplots(figsize=(4,8),constrained_layout=True)
randomsixteenthim=randomsixteenthax.pcolormesh(Shuffledsixteenth,**pc_kwargs)
randomsixteenthfig.colorbar(randomsixteenthim,ax=randomsixteenthax,shrink=0.6)
plt.tick_params(left="off",labelleft="off")
randomsixteenthax.set_xticks(np.arange(len(ticklabels)))
randomsixteenthax.set_xticklabels(ticklabels)
plt.setp(randomsixteenthax.get_xticklabels(),ha="left")
randomsixteenthfig.savefig("FINAL VERSION Random order 0.0625 uM in situ heatmap.pdf",transparent=True)

randomeighthfig,randomeighthax=plt.subplots(figsize=(4,8),constrained_layout=True)
randomeighthim=randomeighthax.pcolormesh(Shuffledeighth,**pc_kwargs)
randomeighthfig.colorbar(randomeighthim,ax=randomeighthax,shrink=0.6)
plt.tick_params(left="off",labelleft="off")
randomeighthax.set_xticks(np.arange(len(ticklabels)))
randomeighthax.set_xticklabels(ticklabels)
plt.setp(randomeighthax.get_xticklabels(),ha="left")
randomeighthfig.savefig("FINAL VERSION Random order 0.125 uM in situ heatmap.pdf",transparent=True)

randomfourthfig,randomfourthax=plt.subplots(figsize=(4,8),constrained_layout=True)
randomfourthim=randomfourthax.pcolormesh(Shuffledfourth,**pc_kwargs)
randomfourthfig.colorbar(randomfourthim,ax=randomfourthax,shrink=0.6)
plt.tick_params(left="off",labelleft="off")
randomfourthax.set_xticks(np.arange(len(ticklabels)))
randomfourthax.set_xticklabels(ticklabels)
plt.setp(randomfourthax.get_xticklabels(),ha="left")
randomfourthfig.savefig("FINAL VERSION Random order 0.25 uM in situ heatmap.pdf",transparent=True)

randomhalffig,randomhalfax=plt.subplots(figsize=(4,8),constrained_layout=True)
randomhalfim=randomhalfax.pcolormesh(Shuffledhalf,**pc_kwargs)
randomhalffig.colorbar(randomhalfim,ax=randomhalfax,shrink=0.6)
plt.tick_params(left="off",labelleft="off")
randomhalfax.set_xticks(np.arange(len(ticklabels)))
randomhalfax.set_xticklabels(ticklabels)
plt.setp(randomhalfax.get_xticklabels(),ha="left")
randomhalffig.savefig("FINAL VERSION Random order 0.5 uM in situ heatmap.pdf",transparent=True)

randomonefig,randomoneax=plt.subplots(figsize=(4,8),constrained_layout=True)
randomoneim=randomoneax.pcolormesh(Shuffledone,**pc_kwargs)
randomonefig.colorbar(randomoneim,ax=randomoneax,shrink=0.6)
plt.tick_params(left="off",labelleft="off")
randomoneax.set_xticks(np.arange(len(ticklabels)))
randomoneax.set_xticklabels(ticklabels)
plt.setp(randomoneax.get_xticklabels(),ha="left")
randomonefig.savefig("FINAL VERSION Random order 1 uM in situ heatmap.pdf",transparent=True)
```
