## Supplemental Appendix S3 for "*Ciona Brachyury* proximal and distal enhancers have different FGF dose-response relationships": readme.txt

Data:

SEVData.csv: Data table containing all of the whole-embryo Summed Enhancer Value measurements for the three reporter constructs

Cell Measurements.csv: Data table containing all of the individual cell reporter expression measurements

InSituForPython.csv: Data table containing all of the semi-quantitative  Bra in situ scores

InSituNonUniform.csv: Data table containing the Bra in situ scores for just the embryos with nonuniform expression


Scripts:

sum_filtering.py: scripts used to extract whole-embryo expression values from confocal stacks.

Fig 2 and Associated Supplement.py: Scripts used for statistical tests and data visualization for figure 2

Fig 3 and Associated Supplement.py: Scripts used for statistical tests and data visualization for figure 3

Fig 4 and Associated Supplement.py: Scripts used for statistical tests and data visualization for figure 4

(file paths are hard coded in these scripts and would need to be adjusted to run on other computers)
