## Supplemental Appendix S3 for "*Ciona Brachyury* proximal and distal enhancers have different FGF dose-response relationships": sum_filtering.py

```
import numpy as np
from veeman_lab_scripts.czifile import CziFile
from os import listdir
from skimage.io import imread
from veeman_lab_scripts.filter import channel_slicer
from tifffile import imsave
from scipy.interpolate import UnivariateSpline
from mahotas import label
import mahotas
import pandas as pd
from scipy.ndimage.filters import median_filter

def name_extractor(image_name):
    embryo_start = image_name.find('embryo')
    embryo_number = image_name[embryo_start + 7:-4]
    if image_name.find('_') == -1:
        embryo_stage = image_name[:image_name.find(' ')]
        enhancer_construct = image_name[len(embryo_stage) + 1 : embryo_start - 1]
        perturbation_name = 'None'

        return perturbation_name, embryo_stage, enhancer_construct, embryo_number

    else:
        perturbation_index = image_name.find('_')
        perturbation_name = image_name[:perturbation_index]
        embryo_stage = image_name[perturbation_index + 1: image_name.find(' ')]
        enhancer_construct = image_name[len(perturbation_name)+ len(embryo_stage) + 2 : embryo_start - 1]

        return perturbation_name, embryo_stage, enhancer_construct, embryo_number


def directory_automatic_projections():
    czi_list = sorted([x for x in listdir() if x[-3:] == 'czi'])

    for image in czi_list:
        #reading in CZI file
        czi_file = CziFile(image)

        #slicing each stack into its seperate channels
        enh_stack = channel_slicer(czi_file, 1)
        phall_stack = channel_slicer(czi_file, 2)

        #sum of slices on phalloidin and enhancer stacks
        summed_phall = np.ndarray.sum(phall_stack, 2)
        summed_enh = np.ndarray.sum(enh_stack, 2)

        #saving output tiffs
        imsave('{}_summed_phall.tif'.format(image[:-4]),
            np.swapaxes(summed_phall, 0, 1))
        imsave('{}_summed_enhancer.tif'.format(image[:-4]),
            np.swapaxes(summed_enh, 0, 1))

def directory_automatic_sums(background_value = 5):
    czi_list = sorted([x for x in listdir() if x[-3:] == 'czi'])
    results = []

    for image in czi_list:
        #reading in CZI file
        czi_file = CziFile(image)

        #slicing each stack into its seperate channels
        enh_stack = channel_slicer(czi_file, 1)
        phall_stack = channel_slicer(czi_file, 2)

        #sum of slices on phalloidin stack
        summed_phall = np.ndarray.sum(phall_stack, 2)

        #finding inflection point of phalloidin values to threshold
        values = np.ndarray.tolist(np.ndarray.flatten(summed_phall))
        values.sort()
        x_range = [x for x in range(len(values))]
        y_spl = UnivariateSpline(x_range,values,s=len(x_range),k=5)
        y_spl_2d = y_spl.derivative(n=2)
        inflection_point = np.where(
            y_spl_2d(x_range[:150000]) == np.amax(y_spl_2d(x_range[:150000])))[0][0]
        threshold_value = values[inflection_point-3000]

        #thresholding phalloidin projection
        summed_phall[summed_phall < threshold_value] = 0

        #labeling thresholded phalloidin image
        labeled = label(summed_phall)[0]

        #finding largest label to get embryo mask
        sizes = mahotas.labeled.labeled_size(labeled)
        embryo_index = np.where(sizes == np.amax(sizes[1:]))[0][0]
        labeled[labeled != embryo_index] = 0
        labeled[labeled == embryo_index] = 1
        counts = np.unique(labeled, return_counts = True)
        mask_size = counts[1][1]

        #summing values of enhancer channel and applying mask
        enh_stack = median_filter(enh_stack, size = 2)
        enh_stack[enh_stack < background_value] = 0
        subtraction_array = np.zeros(enh_stack.shape) - background_value
        subtraction_array = np.where(enh_stack >= background_value, subtraction_array, 0)
        enh_stack = enh_stack + subtraction_array
        summed_enh = np.ndarray.sum(enh_stack, 2)
        enh_sum = np.sum(summed_enh[labeled == 1])

        #so that an image can be written out
        summed_phall = np.ndarray.sum(phall_stack, 2)

        #saving out images
        imsave('{}_embryo_mask.tif'.format(image[:-4]),
            np.swapaxes(labeled, 0, 1).astype(np.uint32))
        imsave('{}_summed_enhancer.tif'.format(image[:-4]),
            np.swapaxes(summed_enh, 0, 1).astype(np.uint32))
        imsave('{}_summed_phall.tif'.format(image[:-4]),
            np.swapaxes(summed_phall, 0, 1).astype(np.uint32))

        #saving values to lists
        results.append(list(name_extractor(image))+ [mask_size, enh_sum])


    output_data = pd.DataFrame(
        results,
        columns = ['perturbation_name', 'embryo_stage', 'enhancer_construct', 'embryo_number',
            'embryo_mask_size', 'summed_enhancer_value'])

    output_data.to_csv('results.csv')


def values_from_stacks(embryo_mask, summed_enh):
    enh_data = imread(summed_enh)
    mask = imread(embryo_mask)
    enh_sum = np.sum(enh_data[mask == 1])
    counts = np.unique(mask, return_counts = True)
    mask_size = counts[1][1]

    return(mask_size, enh_sum)
```
